# The genome of a globally invasive passerine, the common myna (*Acridotheres tristis*)

**DOI:** 10.1101/2023.08.22.554353

**Authors:** Katarina C. Stuart, Rebecca N. Johnson, Richard Major, Kamolphat Atsawawaranunt, Kyle M. Ewart, Lee A. Rollins, Anna W. Santure, Annabel Whibley

## Abstract

In an era of global climate change and massive environmental disturbance, biodiversity conservation is receiving increased attention. Conservation efforts are being greatly aided by genetic tools and approaches, which seek to understand patterns of genetic diversity and how they impact species health and ability to persist under future climate regimes. Invasive species offer vital model systems in which to investigate questions around adaptive potential, with a particular focus on how changes in genetic diversity and effective population size interact with the novel selection regime of the invaded range to drive rapid evolution. The common myna (*Acridotheres tristis*) is a globally invasive passerine, which has undergone multiple concurrent and sequential bottlenecks across its globally invasive range, and yet has established itself across a diverse array of ecological conditions. It is therefore an excellent model species for research both into the persistence of low-diversity populations and the mechanics of biological invasion. To underpin research on the invasion genetics of this species, we present the genome assembly of the common myna, assembled using a backbone of Oxford Nanopore Technologies long reads, alongside an RNA-seq based transcriptome and genome annotation. To provide genomic context for future studies, we describe the genomic landscape of this species, including genome wide allelic diversity, methylation, repeats, and recombination rate, as well as an examination of gene family expansions and contractions. Finally, we use demographic analysis to identify that some native regions underwent a dramatic population increase between the two most recent periods of glaciation, but also reveal artefactual impacts of genetic bottlenecks on demographic analysis.

## 1 INTRODUCTION

Invasive species are organisms that successfully establish outside their native range and expand demographically and spatially, and may negatively impact the new environment, for example causing ecological, environmental, or economic damage (Lockwood *et al*. 2013; Matheson & McGaughran 2022). Invasive species may undergo rapid evolution following introduction to a novel environment, where there may exist a radically different set of biotic and/or abiotic conditions (Thompson 1998; Whitney & Gabler 2008; Buswell *et al*. 2011). Introductions of small numbers of individuals often results in a genetic bottleneck (a sudden reduction in genetic diversity). These features of introduction provide an interesting avenue through which to investigate theories and concepts around rapid adaption in the context of low genetic diversity, genetic inbreeding, mutational load, and novel genetic variation (Schrieber & Lachmuth 2017). Although the increasing affordability of genetic sequencing offers opportunity to assess the mechanisms of rapid adaptation, genomic data is missing for most invasive species (Matheson & McGaughran 2022). Additionally, studying evolutionary processes within invasive populations may be useful for both invasive species management (Clements & Ditommaso 2011; Sillero *et al*. 2020) and the management of vulnerable and/or declining native populations (Willi *et al*. 2022).

*Sturnidae* are a songbird family, comprising more than 100 species spanning a wide range of habitats across much of the globe. Included in this number is the invasive *Acridotheres tristis,* the common myna. The common myna is listed by the IUCN as one of only three bird species on the ‘World’s 100 worst’ invasive species list (Lowe *et al*. 2000). The common myna is characterised by a black head and brown body, with white underwing coverts and a distinctive yellow beak and yellow patch behind the eyes (Fig. 1a). The species is native within central to southeast Asia, and has been either deliberately or accidently introduced to Australia, New Zealand, Israel, South Africa and the United States of America, as well as numerous smaller islands and island groups (e.g. Mauritius, Réunion, Fiji, New Caledonia, Hong Kong, Cayman Islands, Seychelles) (Fig. 1b) (Holzapfel *et al*. 2006; Magory Cohen *et al*. 2019; Beesley *et al*. 2023). The common myna’s ecological impacts are particularly pronounced in island ecosystems (Hughes *et al*. 2017; Feare *et al*. 2022) and have also been studied within Australia, New Zealand, and South Africa, where their widespread geographic coverage has elicited research interest into their invasion history and impacts (Peacock *et al*. 2007; Ewart *et al*. 2019; Beesley *et al*. 2023). The common myna has been the focus of ecological and conservation research owing to their aggressive territorial behaviour that may exclude native avians from nesting structures and foraging areas (Tindall *et al*. 2007; Grarock *et al*. 2012; Rogers *et al*. 2020), though evidence for this is mixed in some environments (Lowe *et al*. 2011). In addition to this, there are also concerns about this species regarding nesting nuisance and crop interference, all of which has led them to be categorised as an agricultural pest in most of their introduced ranges (Tracey & Saunders 2003; Koopman & Pitt 2007).

**Figure 1.**
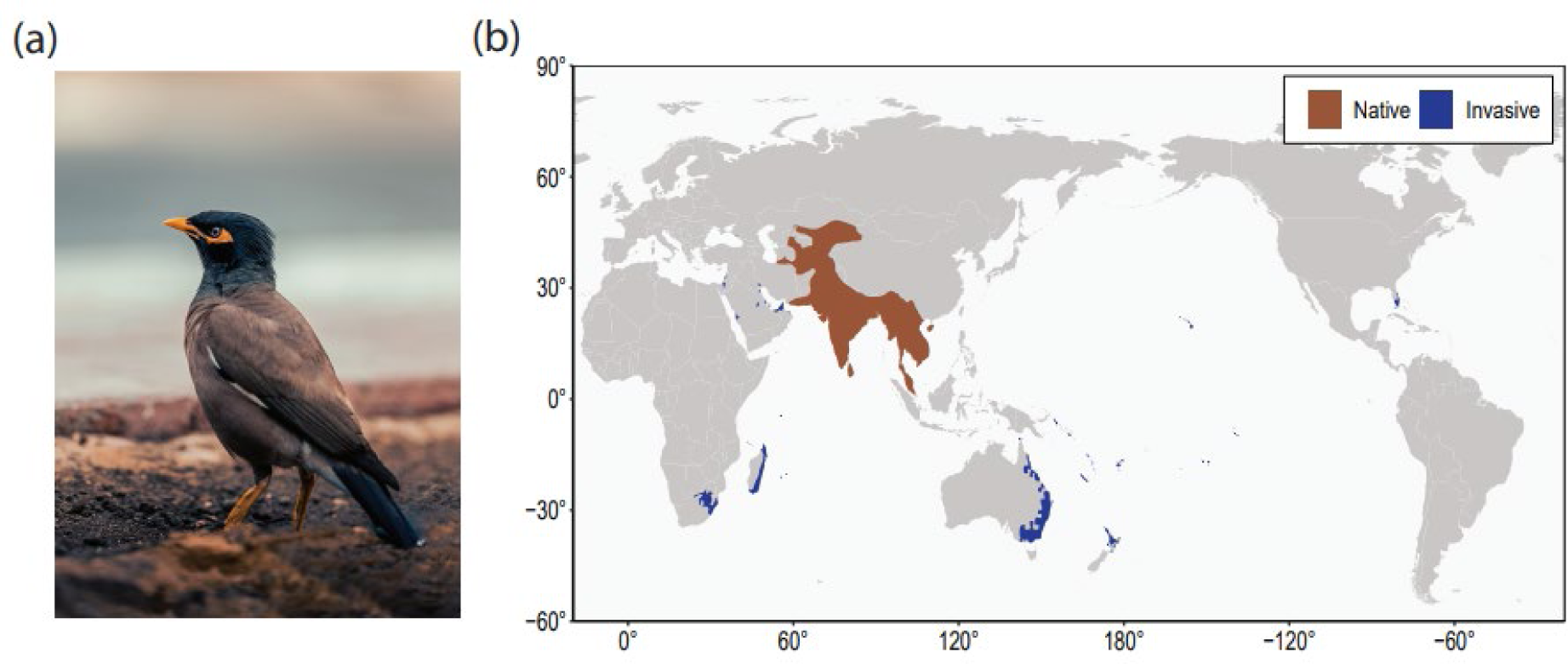
The common myna (*Acridotheres tristis*), and its global distribution across native and invasive ranges. Panel (a) *Acridotheres tristis* (photo credit Rameez Remy). Panel (b) depicts the global distribution of the species, with native range indicated in brown, invasive range indicated in blue.

Evolutionary genomics studies of the common myna are relevant for understanding the impacts of reduced genetic diversity on adaptive potential. This species has undergone multiple concurrent and sequential bottlenecks across its globally invasive range, and yet has established itself across a diverse array of environments (Peacock *et al*. 2007; Magory Cohen *et al*. 2021; Atsawawaranunt *et al*. 2023). Thus, this system gives us the opportunity to study the factors underlying invasion success, rapid adaptation, and population persistence. Further, the common myna is a relative of the European starling (*Sturnus vulgaris*), another globally invasive passerine with a similar introduction history (Stuart *et al*. 2023a). Such interspecies comparisons will provide an opportunity to examine whether molecular evolutionary processes behave in a conserved or stochastic manner across invasions of similar phylogenetic, ecological, and historical nature.

In this manuscript we present the genome assembly of the common myna *A. tristis,* assembled through a combination of long read (Oxford Nanopore Technologies) and linked-read (10x Chromium) sequencing. This reference genome will aid further research into the population genetics and evolutionary genomics of this ecologically important species. To provide genomic context for these studies, we describe the genomic landscape of this species, including recombination, methylation, and single nucleotide polymorphisms (SNPs), as well as briefly examining gene family expansions and the demographic history of native and invasive populations.

## 2 MATERIALS AND METHODS

### 2.1 Sampling and sequencing

A male common myna was collected from Newcastle, Australia (-32.935, 151.751) on 15/07/2014 and the snap-frozen tissues (liver, heart, testis, breast muscle) were stored at -80°C (Australian Museum Registration Number O.76569, local ID 13099). Extractions (DNA and messenger RNA) was conducted, and 10x Chromium linked-read, Oxford Nanopore Technologies long-read, and short read cDNA sequencing was performed (Appendix 1: Sampling, extraction, and sequencing).

Additionally, representative common myna individuals from globally distributed native and invasive populations (Ewart *et al*. 2019; Atsawawaranunt *et al*. 2023) were whole genome resequenced (WGR) using short-read whole genome resequencing on the Illumina Novaseq platform (150 bp paired end reads) with sequencing brokered by Custom Science, Australasia (Table S1).

### 2.3 Genome assembly

The assembly process is summarised in Fig. 2. First, we processed raw ONT (Oxford Nanopore Technologies) long-reads with GUPPY v6.2.1 (for config settings see: Table S2). After basecalling, PORECHOP v0.2.4 was used to detect and remove residual sequencing adaptors (Wick *et al*. 2017). These reads were assembled into an initial assembly with FLYE v2.9.1 (Kolmogorov *et al*. 2019) using the settings ‘--no-alt-contigs’ and ‘--scaffold’. The ONT reads were mapped back to the draft assembly and polished using MEDAKA v1.4.3 (Oxford Nanopore Technologies 2018) with default parameters and the r941_min_sup_g507 model. The assembly was then further polished through two iterations of NEXTPOLISH v1.4.1 (Hu *et al*. 2020), using Chromium 10x reads that had been stripped of their barcodes using SCAFF10X v5.0 (https://github.com/wtsi-hpag/Scaff10X) and then quality filtered using TRIMGALORE v0.6.7 (Krueger 2021) with default parameters plus the “–2-colour” flag enabled. Reads were mapped to the draft genome using bwa mem v0.717 and alignments were sorted and compressed using SAMTOOLS v1.14 (Danecek *et al*. 2021). The linked-read data was not used for scaffolding as it did not improve assembly continuity or provide useful long-range information. The polished genome was then manually curated to correct for misassembles (Appendix 2: Manual genome curation). Finally, the assembly was then aligned to the Vertebrate Genomes Project (VGP) common myna genome (GCA_027559615.1) using RAGTAG v2.1.0 (Alonge *et al*. 2022) to produce synteny based superscaffolds (one VGP contig was excluded from scaffolding, see: Appendix 3, Fig. S4, Fig. S5). These scaffolds were then renamed based on their synteny with the major chromosomes of the zebra finch genome (Fig. S6). The final genome assembly produced from the above protocol we refer to as AcTris_vAus2.0.

**Figure 2.**
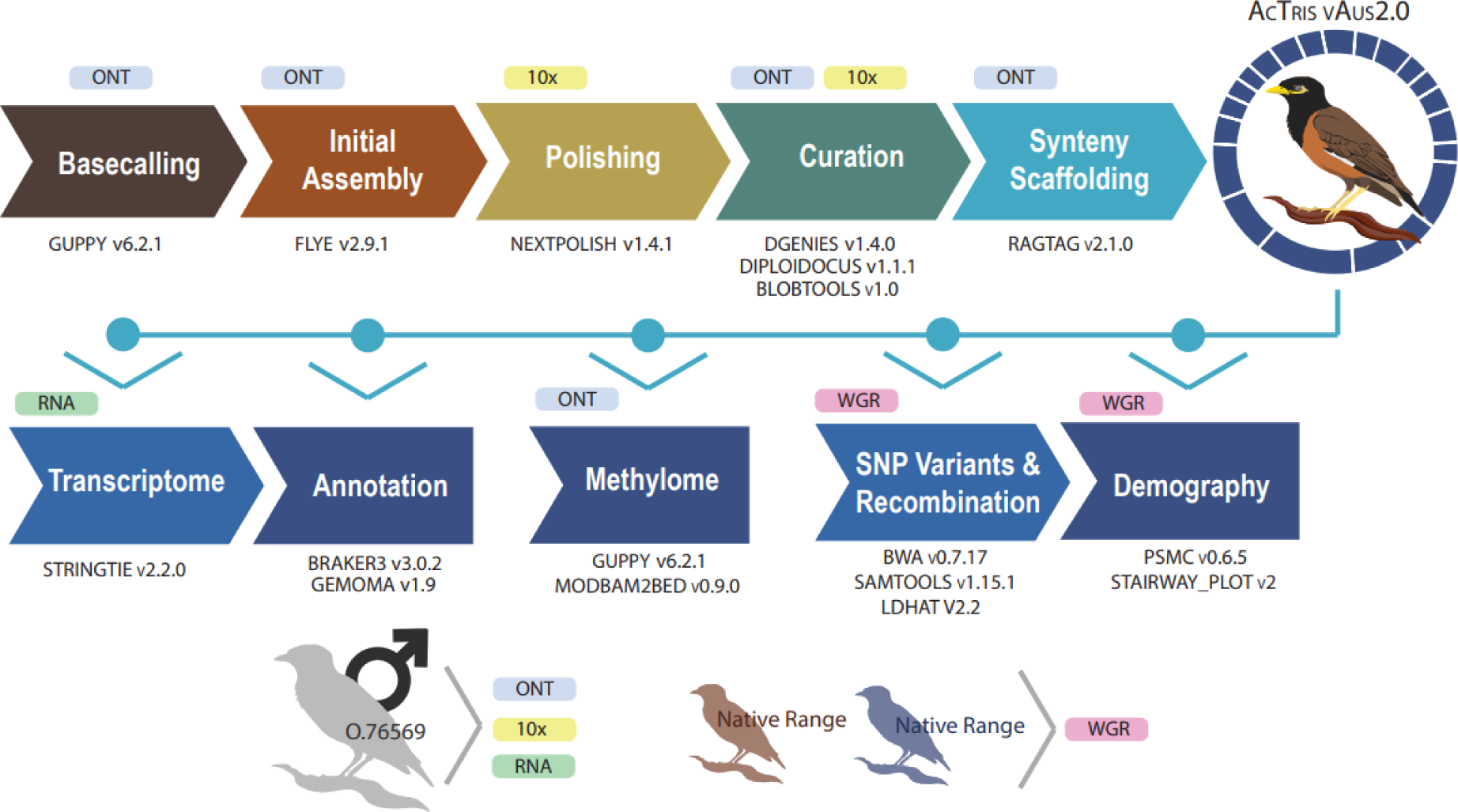
The common myna (*Acridotheres tristis*) genome workflow summary information. ONT = Oxford Nanopore Technologies long reads. 10x = 10x chromium linked Illumina reads. RNA = Illumina short reads. WGR = Illumina whole genome resequencing.

### 2.4 Assembly evaluation

We used k-mer frequency analysis through JELLYFISH v2.3.0 to assess genome size. A k-mer histogram was produced using all the trimmed linked read gDNA raw data for an initial value of 20-mer based on an approximated genome size of just above 1 Gb (k-mer values 18, 19, 21, and 22 were also assessed). Counts for k-mer values of 7 or below were attributed to extremely rare reads or sequencing errors, and were removed. The genome size was then estimated by dividing the total number of k-mers over all k values by the mean coverage.

In addition to this, MERQURY v1.3 (Rhie *et al*. 2020) was used to assess completeness and the assembly consensus quality value (QV) (using a genome size of 1.040e9, a k-mer value of 20 was selected to construct a MERYL v1.4 database).

We assessed the AcTris_vAus2.0 genome (as well as the VGP common myna genome) assembly contiguity with SEQSUITE v1.27.0, and completeness with BUSCO v5.3.2 (Simão *et al*. 2015) using genome mode and ‘Aves’ lineage from the ODB10 dataset.

### 2.5 Transcriptome assembly

We assembled a genome-guided transcriptome assembly from the three Illumina-sequenced RNA tissues (liver, heart, testes). Raw reads were trimmed and quality filtered using FASTP v0.23.2 (Chen *et al*. 2018) (default settings), before being mapped to AcTris_vAus2.0 using HISAT2 v2.2.1 (Kim *et al*. 2019) (settings: --rna-strandness FR –dta --phred33). The resulting sam files were sorted and converted to bam using SAMTOOLS v1.15.1, before being assembled into a gtf file using STRINGTIE v2.2.0 (Kovaka *et al*. 2019). Each tissue was initially assembled into individual gtf files, and these tissue specific transcriptomes were then combined into one using the STRINGTIE –merge function. The overlap across tissues within this merged transcriptome was assessed using GFFCOMPARE v0.12.6 (Pertea & Pertea 2020). We assessed the completeness of this three-tissue transcriptome using BUSCO (--mode transcriptome).

### 2.6 Genome annotation

For the genome annotation of the common myna, we generated ab-initio gene predictions using BRAKER v3.0.2 (Hoff *et al*. 2016; Brůna *et al*. 2021), homology based gene predictions using GEMOMA v1.9 (Keilwagen *et al*. 2018), and merged these into a singular annotation using TSERBA v1.1.0 (Gabriel *et al*. 2021). First, we soft masked the genome using the joint repeat library described below (see section 2.7.3: Transposable Element and Repeat Annotation). BRAKER was provided with the three separate RNAseq data bam files (one for each tissue, see 2.5: Transcriptome assembly), and the UniProt/Swiss-Prot database (UniProt Consortium 2019) as evidence. BRAKER employs both RNA and protein evidence to run GENEMARK-EPT (Brůna *et al*. 2023), PROTHINT (Brůna *et al*. 2020), and train AUGUSTUS, with redundant training gene structures filtered out using DIAMOND (Buchfink *et al*. 2015). GEMOMA was run concurrently to this, and was provided the annotation information for 26 avian species (Table S3) (parameters: tblastn=false GEMOMA.m=200000 GEMOMA.Score=ReAlign AnnotationFinalizer.r=SIMPLE pc=true o=true).

These two annotations were then merged using TSEBRA, with the BRAKER gene set enforced so that the final annotation would retain the species specific ab-initio sequences predicted by BRAKER, but would merge in the homology based sequences called by GEMOMA (configuration file settings: hint weightings P 0.1, E 10, C 5, M 1; intron_support 0.1, stasto_support 1; e_1 0.1, e_2 0.5, e_3 0.05, e_4 0.18). We generated functional annotation of protein-coding genes using EGGNOG-MAPPER v2.1.10 (Cantalapiedra *et al*. 2021) using DIAMOND (-m diamond) to perform the protein sequence searches.

We assessed the completeness of the final annotation (as well as the individual BRAKER and GEMOMA annotations) using BUSCO (--mode transcriptome). We assessed the quality of predicted transcripts using SAAGA v0.7.7 (Stuart *et al*. 2022a) with the Ensembl *Gallus gallus* proteome (GCF_016699485.2) as a reference, and summarised these statistics overall as well as separately for macro and micro chromosomes, where we define macrochromosomes as chromosomes 1, 1A, 2, 3, 4 and 5 (Fig. S7). We generated annotation statistic summaries using the AGAT (Dainat 2020) agat_sp_functional_statistics.pl script, and used BEDTOOLS to plot gene density in 1 Mb windows across the whole genome.

### 2.7 Genomic landscape

We explored the landscape of SNPs, methylation, repeat and transposable element (TE) content, and linkage disequilibrium (LD) based inference of recombination along the genome assembly of AcTris_vAus2.0, to provide context and resources for future genomic studies on this species.

#### 2.7.1 SNP variant density

The WGR data from 15 individuals across two native range sample sites (Tamil Nadu = TN and Madhya Pradesh = MP, Table S1) were used to quantify genome wide SNP density. Raw reads were processed using TRIM_GALORE v0.6.7 (Krueger 2021) and the reads were mapped to the AcTris_vAus2.0 genome assembly using BWA v0.7.17 (Li 2013) *mem*, before being processed by SAMTOOLS into sorted BAM files. Duplicate reads were marked using PICARD v2.26.10 (‘Picard toolkit’ 2019) *MarkDuplicates*, and variants jointly called across samples using BCFTOOLS v1.13 (Danecek *et al*. 2021) *mpileup* (-C 50 -q 20 -Q 25), *call* and *view* functions. Indels were excluded from the dataset, and SNPs were filtered for a minimum depth of 5, a maximum depth of 50, and non-variant sites were removed using VCFTOOLS v0.1.15 (Danecek *et al*. 2011) (--mac 1). SNP density was then calculated for 1 Mb bins along the genome, and was also summarised across macro and micro chromosomes.

#### 2.7.2 Methylation profiling

We used GUPPY v6.2.1 to perform extended methylation base calling of the ONT long-reads against the AcTris_vAus2.0 genome, with each flow cell batch run separately (for config files see: Table S2). Reads that were assigned a ‘pass’ score were then combined and sorted using SAMTOOLS, before we aggregated modified base counts using MODBAM2BED v0.9.0 (https://github.com/epi2me-labs/modbam2bed) (-m 5mC) to identify 5-methylcytosine (5mC) in a CpG context. BEDTOOLS *coverage* was then used to assess this genome methylation coverage across three flow cell types separately in 1 Mb windows to check for consensus (Fig. S8), before the bed files of all three flow cell types were combined into a joint MODBAM2BED run. Information about methylated read proportion for each methylated CpG site (with a minimum read depth of 5) was calculated, and CpG site counts were calculated for 1 Mb windows along the genome using BEDTOOLS *coverage* (this was done separately for all CpG sites, and those with 75% or more methylated reads). The individual sites of methylation (filtered for a minimum coverage of 5) along the genome were then merged into methylated regions using DMRFINDER v0.3 (Gaspar & Hart 2017) using default settings other than a minimum count number of 5 (-r 5) and a minimum CpG sites per region set to 15 (-c 15) because we chose to focus only on those windows with a high density of CpGs. The proportion of methylated reads in each region were then summarised across macro and micro chromosomes, up to 5kb upstream of gene sequences, and within transposable elements.

#### 2.7.3 Repeat and transposable element annotation

A repeat library was generated for the AcTris_vAus2.0 assembly using several means. We first generated a species-specific repeat library following the MAKER2 advanced repeat library construction protocol (http://weatherby.genetics.utah.edu/MAKER/wiki/index.php/Repeat_Library_Construction-Advanced) with miniature inverted-repeat transposable elements identified using MITE-TRACKER (Crescente *et al*. 2018). We also identified TEs in the genome using EARLGREY v2.0 (Baril *et al*. 2022), which employs REPEATMASKER v4.1.2 (Smit *et al*. 2013), REPEATMODELER v2.0.2 (Flynn *et al*. 2020) and the Dfam 3.6 database (Storer *et al*. 2021) in a fully-automated TE annotation pipeline to create de novo consensus sequences. Then, we assessed the repeat content of the AcTris_vAus2.0 genome using REPEATMASKER, using a joint repeat library by combining the MAKER2 advanced repeat library, EARLGREY consensus sequences, and the Aves lineage specific sequences from the RepeatMasker repeat database. We also ran the VGP common myna genome through EARLGREY and annotated repeats using REPEATMASKER (using the same repeat library generated above but with the genome assembly specific EARLGREY library substituted in).

#### 2.7.4 Recombination profile

To generate a linear recombination landscape for each of the common myna superscaffolds, we used the SNP data generated above (see 2.7.1: SNP variant density) to run LDHAT V2.2 (McVean *et al*. 2004), which can estimate recombination rates from linkage disequilibrium measures for SNPs in population genetic data. We analysed the two separate sample sites together as we believe the utility of increasing sample size to help resolve broad recombination patterns along chromosomes outweighs possible impacts of pooling individuals from two separate geographic sample sites. The SNP variant file was split into each separate chromosome, as LDHAT must be run separately for each scaffold, and converted into LDHAT format using VCFTOOLS. We did not perform minor allele frequency filtering, because previous research has shown removing singletons does not have a large impact on program output when high-quality reference genomes are used (Stukenbrock & Dutheil 2018). However, we did use VCFTOOLS to thin the SNP data (--thin 1,000) to reduce computation time and resources. Each chromosome was then run through LDHAT *interval* (-its 10000000 -bpen 5 -samp 5000) and then LDHAT *stats* (-burn 250) to summarise the output. The population-scaled recombination rate estimate Rho (ρ = 4N_e_r, where r is the recombination rate) was then plotted along each scaffold’s length to illustrate the linear landscape of recombination along each chromosome’s length. We also examined linkage disequilibrium decay in a pairwise manner between SNPs on the same chromosome (up to 10 Mb away from each other) using VCFTOOLS using the “--geno-r2” function (--ld-window-bp 10,000,000).

### 2.8 Gene family expansions

We identified gene families and orthologous gene clusters of 12 species and identified expansions within these using ORTHOFINDER v2.5.2 (Emms & Kelly 2019) and CAFE5 (Mendes *et al*. 2020). We used two outgroup species of *Homo sapiens* (GCF_000001405.26) and *Mus musculus* (GCF_000001635.2), and nine ingroup bird species of *Lonchura striata domestica* (GCF_002197715.1), *Taeniopygia guttata* (GCA_003957565.1), *Gallus gallus* (GCF_016699485.2), *Ficedula albicollis* (GCF_000247815.1), *Cyanistes caeruleus* (GCF_002901205.1), *Serinus canaria* (GCF_000534875.1), *Parus major* (GCF_001522545.2), *Zonotrichia albicollis* (GCF_000385455.1) and *Sturnus vulgaris* (vAU1.0 from Stuart *et al*. 2022), alongside the AcTris_vAus2.0 genome annotation. The longest transcript for each gene in the annotation file for each species was identified using AGAT agat_sp_keep_longest_isoform.pl and the protein sequence extracted using the matching genome assembly and GFFREAD v0.12.7 (Pertea & Pertea 2020). We used SEQKIT v2.4 to remove protein sequences less than 30 amino acids long. In addition, a small number of species had translated protein sequences with premature stop codons; these were filtered out with SEQKIT. We ran these final protein sequences through ORTHOFINDER (-M msa -S blast -I 1.3) to produce a species tree and identify phylogenetic hierarchical orthogroups (HOGs) that had expanded or contracted on specific branches. Timetree (Kumar *et al*. 2022) was then used to make the species tree ultrametric, which was then used alongside a summary of HOGs across the 12 species as input for CAFE5 to determine which of these HOG had significant expansions or contractions in taxa and linages. We ran CAFE5 five times with gamma rate categories count (-k) set to 1, 2, 3, 4 and 5 respectively. The log files of each run were checked for convergence, and the run with the smallest likelihood score was retained (k=4). We identified which HOGs had reported a significant contraction or expansion across the phylogeny (p-value < 0.05), and filtered these for HOGs with expansions or contractions identified in AcTris_vAus2.0. These HOGs were then mapped to their corresponding orthogroup (OG), and the largest five sequences associated within each OG were functionally annotated with their associated gene ontology (GO) terms using INTERPROSCAN v5.51 (Quevillon *et al*. 2005) (using the –goterms flag). The GO terms biological processes were then summarised separately for expanding and contracting sequences using REVIGO (Supek *et al*. 2011).

### 2.9 Demographic inference from assembly

To estimate effective population size (N_e_) over ancient timescales for the common myna we used the program PSMC v0.6.5 (Li & Durbin 2011), which employs hidden Markov models in a coalescent approach to identify historical recombination events in a single diploid genome. A single whole genome resequenced representative individual was selected from four native range sample sites and six invasive range sample sites (Table S1). The raw reads of these individuals were individually put through a variant calling pipeline similar to that described above (see section 2.7.1: SNP variant density), except that SNPs within 10 bp of an indel or overlapping repeat regions were excluded (see section 2.7.3: Repeat and transposable element annotation) following the PSMC protocol used by Schield *et al*. (2022). We ran PSMC for 30 iterations (-N 30), with the upper limit of time to most recent common ancestor set to 5 (-T 5), an initial h:q value of 5 (-r 5), and free atomic time intervals set to (4+30*2+4+6+10) based on recommendations for avians (Nadachowska-Brzyska *et al*. 2015). We performed bootstrapping (50 iterations) to check for variation in N_e_ estimates. Results were then scaled using an estimated generation time of 2 years (approximate age of first breeding: Feare & Craig 1999) and a yearly mutation rate of 2.3 x 10^-9^ based on related avian species estimates from (Nadachowska-Brzyska *et al*. 2015; Smeds *et al*. 2016).

To understand more recent N_e_ demographic changes, we used the tool STAIRWAY PLOT v2 (Liu & Fu 2020), which uses the population site frequency spectrum (SFS) for inferring demographic history. We ran this analysis first on one of the native range sample sites (Tamil Nadur, India, IND TN N = 8) using the SNP data called above (section 2.7.1: SNP variant density). Then, because we were interested in the effects strong genetic bottleneck effects within invasive ranges have on demographic history, we called and analysed SNP data for an invasive range sample site as well (Leigh, New Zealand, NZ LEI N = 8). SFS information was obtained for variant sites using VCF2SFS in R (Liu *et al*. 2018), and total number of observed nucleic sites (L) was set to the total sites in our SNP data (variant + non-variant) once filtered to only include individuals within each of the sample sites. A generational mutation rate was set to 4.6 x 10^-9^ (the yearly mutation rate above x generation time used in PSMC analysis). All other parameters were run at their default settings.

## 3 RESULTS & DISCUSSION

### 3.1 Genome assembly

A combination of long and short read technologies was used to assemble, scaffold, and polish the genome of the common myna (AcTris_vAus2.0). A total of 23.5 Gb of ONT long reads (Table 1) were used to create an initial genome assembly of 1,648 contigs with a contig N50 of 11.3 Mb and a total length of 1.046 Gb, a continuous assembly given ONT coverage was lower than for many other avian genomes (Rhie *et al*. 2020; Peona *et al*. 2021; Bailey *et al*. 2023). Sequence curation and trimming reduced this down to 597 contigs, with a contig N50 of 10,406,399 and a total length of 1.041 Gb. After species-specific synteny scaffolding, the final genome comprised a total of 256 scaffolds, with a median scaffold length of 3,369 bp and 98.9% of the assembly that could be anchored to putative chromosomes (Table 2, Fig. S7).

**Table 1.**
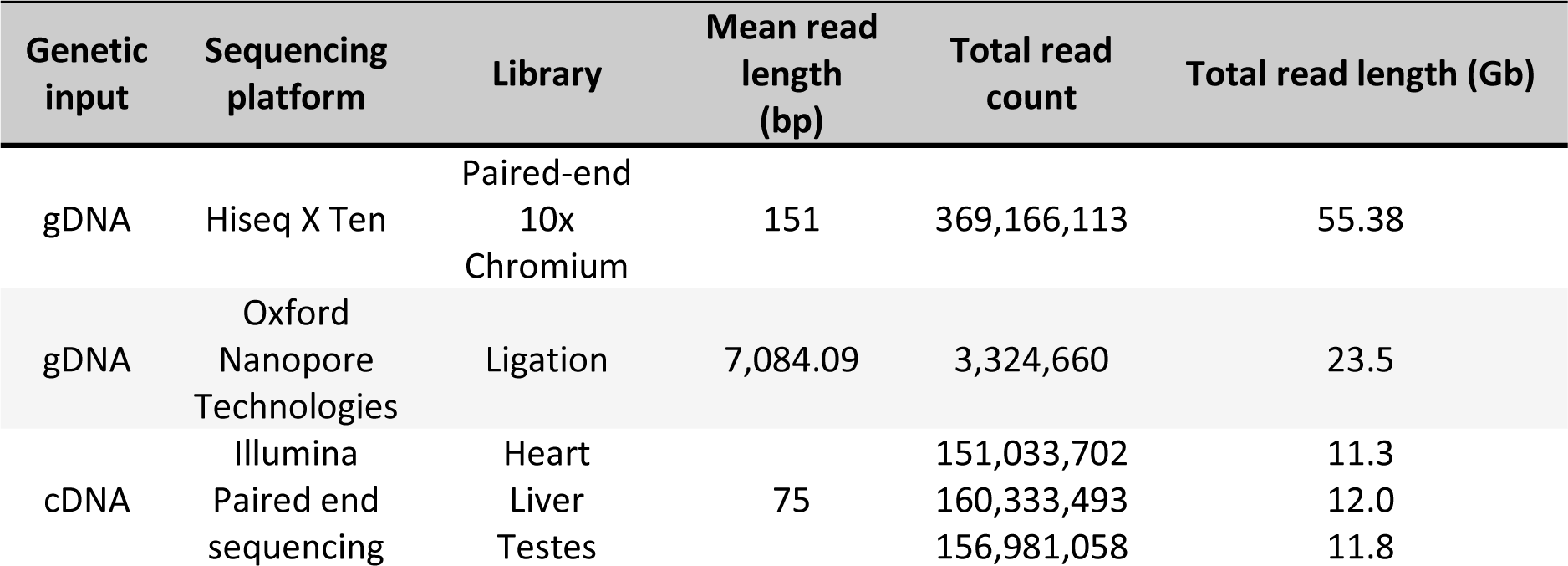
Library information of all sequencing data used in the construction of the *Acridotheres tristis* reference genome and annotation. Total read count, total read length, and median/mean insert size all raw totals before quality filtering. All sequence data was obtained from the same individual O.76569.

**Table 2.** Genome assembly statistics for the *Acridotheres tristis* AcTris_vAus2.0 genome.

| Assembly Statistic | AcTris_vAus2.0 |
| --- | --- |
| Total length (bp) | 1,040,539,946 |
| Number of scaffolds | 256 |
| Scaffold N50 (bp) | 72,486,765 |
| Scaffold L50 | 5 |
| Largest scaffold (bp) | 150,861,042 |
| Mean scaffold length (bp) | 4,064,609.16 |
| Median scaffold length (bp) | 3,369 |
| Number of Contigs | 597 |
| Contig N50 (bp) | 10,406,399 |
| Contig L50 | 30 |
| Gap (N) length (bp) | 34,958 (0.00%) |
| GC content (%) | 41.85% |
| BUSCO (genome: Aves) | 8,338 |
| Complete | 8,108 (97.2%) |
| Complete (single copy) | 8,077 (96.9%) |
| Complete (duplicated) | 31 (0.4%) |
| Fragmented | 44 (0.5%) |
| Missing | 186 (2.2%) |

Analysis of genome completeness using BUSCO indicated that 97.3% of expected single-copy orthologs were complete and single-copy within the genome (Table 2). Genome size for the common myna was estimated to be approximately 1.162 Gb using k-mer analysis (Fig. S9, Table S4) of the 55.38 Gb raw short reads (Table 1), which placed AcTris_vAus2.0 at roughly 90% completeness assuming the kmer estimates were not biased due to fluctuations in sequencing data coverage. Together, this evidence suggests that the remaining length of the genome likely is biased towards non-coding regions (Beauclair *et al*. 2019). This gap between the estimated genome size and final assembly length is not unusual because avian species, even with their fairly small and repeat-light genomes, have been found to be missing 7-42% of their expected genome length in previous genome assemblies (Peona *et al*. 2018). While this is improving as long-read sequencing becomes more commonplace, enabling characterisation of more hard-to-assemble regions (Driver & Balakrishnan 2021; He *et al*. 2021), there is still much about avian genomes left to explore (Botero-Castro *et al*. 2017; Bravo *et al*. 2021; Rhie *et al*. 2021; Peona *et al*. 2022)

For our genome assembly of the common myna, chromosome identities were assigned to superscaffolds based on synteny to the model passerine the zebra finch, given avian karotypes are highly conserved (Sharma *et al*. 1980; O’Connor *et al*. 2019). Our assembly, like many other avian genomes, was missing chromosome 16, which is highly repetitive owing to containing many copies of the MHC gene and hard to assemble (Miller & Taylor 2016). The largest superscaffold length was 151 Mb in length (Table 2, Fig. S7), and equated to the autosomal chromosome 2 which is the largest for most passerine species (though not for all avians; Waters *et al*. 2021), while the major (Z) sex chromosome was the fourth largest of the assembled superscaffolds, which aligns with karyotype information available for this species (Sharma *et al*. 1980).

We compared the overall genome completeness statistics of this version of the common myna genome to the VGP common myna genome. While AcTris_vAus2.0 is shorter than the VGP genome by approximately 150 Mb (Table S5), a majority of this extra length is contained on smaller contigs that are at presently unassigned to a chromosome (Fig. S1) and places the final genome length of 1.193 Gb above the k-mer based genome size estimate for our genome. The two genome versions have comparable BUSCO completeness scores (97.3% for AcTris_vAus2.0, 97.1% for the VGP genome), meaning that likely these additional unplaced contigs are repeat heavy and/or non-protein coding regions, as is typical of genomic regions that are harder to place within larger scaffolds (Weissensteiner & Suh 2019). Additionally, the repeat and TE content of the VGP genome was much higher than that of AcTris_vAus2.0 (18.8% and 9.8% respectively, Fig. S10). With passerine genomes typically containing 10% or less repeat sequence content (Zhang *et al*. 2014; Kapusta & Suh 2017), it is thus likely that some of this extra length contains inappropriately expanded repeats (Peona *et al*. 2021) or haplotypes (Guiglielmoni *et al*. 2021). These differences in genome length and repeat content (and genome structure: Appendix 3) may in part reflect differences in the primary sequencing technologies used in each assembly (PacBio for the VGP genome, and ONT for AcTris_vAus2.0), though will also reflect pipeline decisions downstream of the initial genome assembly and possibly some genuine biological differences between source populations (AcTris_vAus2.0 is an Australian sourced bird, the VGP common myna genome was sourced from Israel). However, we explicitly note that the VGP genome is an early release and will likely undergo further curation. Although we generated the linked read 10x dataset to aid in scaffolding, initial exploration of these datasets revealed that it did not significantly improve assembly contiguity for the genome presented in this study. The VGP genome therefore provided a species-specific means of generating chromosome level superscaffolds from the initial long read assembly in the absence of Hi-C data for AcTris_vAus2.0, demonstrating how valuable research efforts such as the VGP (Rhie *et al*. 2021) and B10k (Zhang 2015) are for enhancing the science of smaller consortiums and lab groups. In the era of genome assemblies being created by large conglomerates *en masse*, there is still an important space for creation and release of additional genome versions of the same species, particularly when achieved through different sequencing technologies. Such efforts are complementary and will ultimately form the backbone of data needed for high quality genome graphs and pan-genomes to better capture structurally diverse regions of the genome (Kim *et al*. 2019; Sherman & Salzberg 2020).

### 3.2 Transcriptome & Annotation

We obtained a total of 35.1 Gb short-read RNA sequencing data over three different tissues (Table 1), from which we generated 32,617 transcript sequences. RNA sequencing of the testes tissue yielded the highest number of unique transcripts, likely because of the alternative splicing occurring in this sex organ (Song *et al*. 2020; Mueller *et al*. 2021), while unique RNA contributions to the transcriptome was near identical for the heart and liver tissues (Fig. S11). BUSCO analysis of the transcriptome revealed that despite just three tissues being combined in its creation, only 14.9% of BUSCO genes were missing from this final transcriptome (Table 3), an amount comparable to other short read transcriptomes (Richardson *et al*. 2017). Nevertheless, expanding tissue diversity in the common myna is an integral next step for full characterisation of the transcriptomic landscape of the species, because a large portion of missing transcripts are thought to be due to tissue specific expression (Yin *et al*. 2019). The transcriptome was then used for gene prediction when completing the annotation of the AcTris_vAus2.0 genome assembly.

**Table 3.** Transcriptome and proteome statistics for the *Acridotheres tristis* AcTris_vAus2.0 genome.

| Statistic |  | AcTris_vAus2.0 |
| --- | --- | --- |
| Transcriptome<sup>a</sup> |  |  |
| BUSCO (transcriptome: Aves) |  | 8,338 |
| Complete |  | 6,917 (83.0%) |
| Complete (single copy) |  | 4,162 (49.9%) |
| Complete (duplicated) |  | 2,755 (33.0%) |
| Fragmented |  | 184 (2.2%) |
| Missing |  | 1,237 (14.8%) |
| Annotation<sup>b,*</sup> |  |  |
| BUSCO (transcriptome: Aves) |  | 8,338 |
| Complete |  | 8,211 (98.5%) |
| Complete (single copy) |  | 8,157 (97.8%) |
| Complete (duplicated) |  | 54 (0.6%) |
| Fragmented |  | 47 (0.6%) |
| Missing |  | 80 (1.0%) |
| Genes | Total number | 19,836 |
|  | Average length | 23,344 bp |
|  | Mean mRNAs per gene | 2.1 |
| mRNA | Total number | 41,104 |
|  | Average length | 32,068 bp |
|  | Mean exons per mRNA | 12.2 |
| CDS | Total number | 41,104 |
|  | Average length | 1,955 |
|  | Average intron in CDS length | 2,680 |
| Exons | Total number | 502,825 |
|  | Mean length | 160 |
| Gene Function | Ontology term | 17,016 |
|  | Protein Family | 16,289 |

<sup>a</sup> Based on three tissue RNA-seq transcriptome assembled in this study
<sup>b</sup> Based on the TSEBRA merged BRAKER and GEMOMA annotation
\* filtered for longest transcript

The final annotation of AcTris_vAus2.0 identified 19,836 gene and 41,104 mRNA sequences across the genome. When restricted to just the longest transcript per gene, the total gene sequence coverage was 31,728,307 bp, equating to 3.05% of the genome’s length (Table 3). The final genome annotation had a BUSCO completeness score of 98.4% (Table 3), and was a merge of two gene models, one produced by GEMOMA and one BRAKER. GEMOMA, being homology based and thus biased towards easier to predict genes in more conserved genomic regions achieved the highest BUSCO scores (97.7%, Table S6), and a total gene and transcript count of 22,216 and 69,476 respectively. Nevertheless, BRAKER performed well, identifying 13,773 gene sequences and 18,261 transcript sequences, with a fairly high BUSCO score (88.0%, Table S6). For the final annotation, EGGNOGG assigned an identity to 17,016 genes (Table 3).

Annotation quality was assessed using SAAGA (Stuart *et al*. 2022a). The 19,836 longest transcript protein sequences were mapped to the high-quality *Gallus gallus* reference proteome (GCF_016699485.2), with 15,212 returning successful hits (76.7%) and 4,624 transcripts returning no hit against this reference (23.3%). Sequences with successful hits were on average longer (600 vs 309 amino acids) and contained more exons (11 vs 4 exons per sequence) compared to sequences failing to match. While longer unknown proteins may be indicative of legitimate novel sequences, it is likely that these also contain short sequences of incorrectly predicted or fragmented gene sequences.

### 3.3 Genomic landscape

We explore the genomic landscape of the common myna genome version AcTris_vAus2.0. Because of their repeat-sparse genomes, short generation times, and diverse ecological interactions, invasive passerines pose promising model systems in which investigate eco-evolutionary processes such as rapid adaptation. Further, the European starling *Sturnus vulgaris* is also listed in the IUCN’s ‘World’s 100 worst’ invasive species list (Lowe *et al*. 2000), and comparative genomic studies using these two species will help examine the predictive or stochastic nature of rapid evolution (Stuart *et al*. 2023b). To this effect, we use our newly constructed genome to characterise patterns along the genome (macro, micro and major sex chromosomes) for important genetic features including single nucleotide polymorphisms (SNPs), gene density, methylation (specifically CpG sites), transposable element (TE) and repeat content and finally linkage disequilibrium (LD) based recombination estimates.

Across the 15 whole genomes used to characterise allelic diversity, we identified a total of 22,992,315 SNPs post filtering (variant sites of minimum depth 5, maximum allele depth 50), which represents 2.2% of the genome. We plotted this whole genome variant data (Fig. 3; track 1) to visualise regions where variant density is high as indicated by peaks, and low trough regions which are indicative of locally reduced variant density and thus may be interpreted as regions of high conservation across conspecifics of the species. Variant density was fairly consistent across the genome, with chromosome ends reporting relatively fewer SNPs likely due to difficulties in mapping to these low-complexity regions (Li 2014). The deficit of SNPs on chromosome Z in the common myna reflects results found in other avian species (Lavretsky *et al*. 2015; Stuart *et al*. 2022b), which is notable given the disproportionate role the major sex chromosome plays in adaptation and speciation (Meisel & Connallon 2013). When variant density was examined across macro and micro chromosomes separately, we observed that macrochromosomes and microchromosomes had very similar variant density profiles (Fig. 4a).

**Figure 3.**
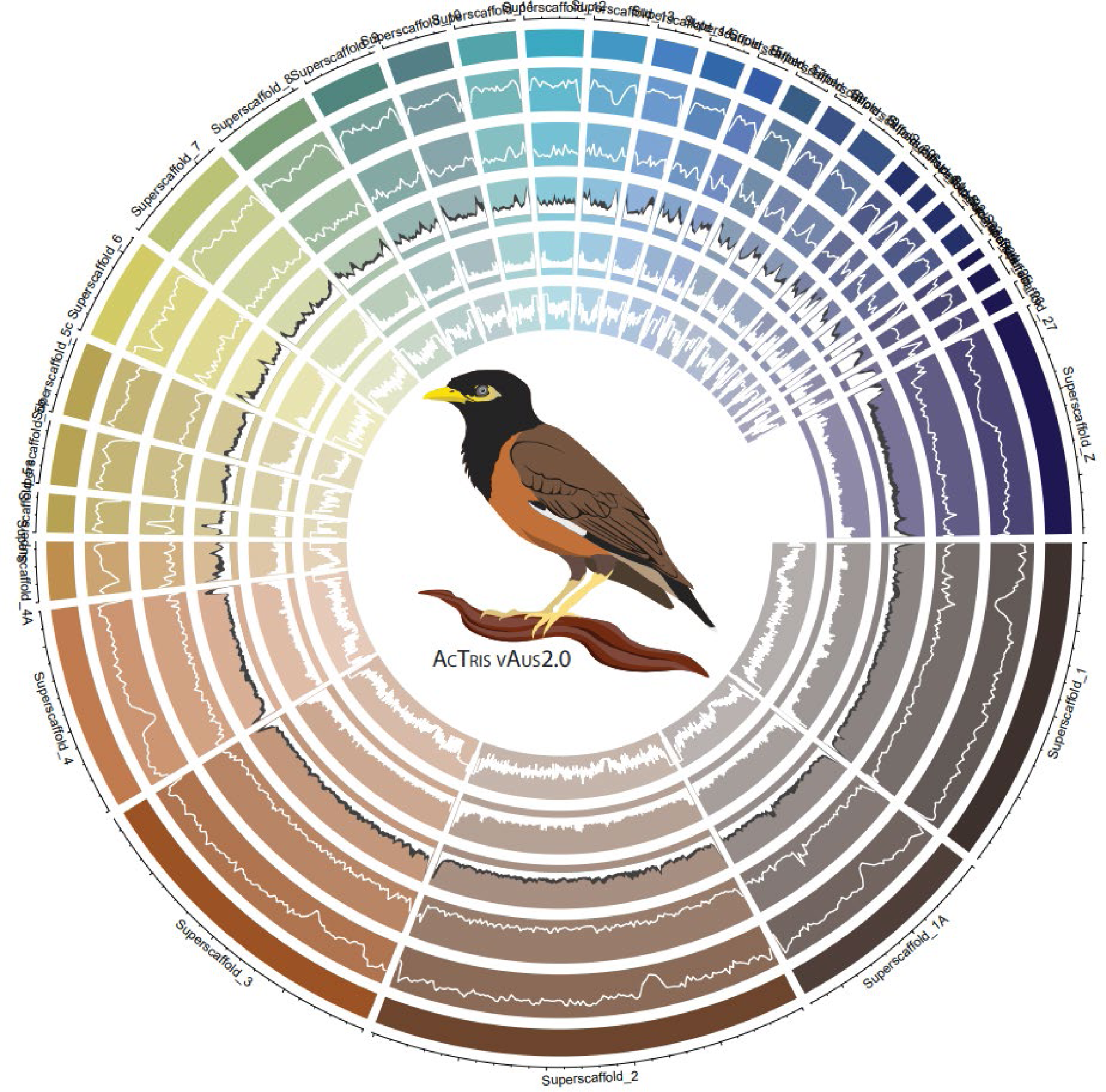
Chromosome coverage plots for the *Acridotheres tristis* AcTris_vAus2.0 genome. CIRCLIZE plot of the 30 largest superscaffolds, with tracks (from the outside in) SNP density (1 Mb windows), gene density (1 Mb windows), CpG site density (1 Mb windows, CpG sites with 0-75% methylated reads in white, and 75-100% methylation in grey), and repeat density (0.5Mb windows), and recombination Rho (log corrected. Not plotted for Z chromosome).

**Figure 4.**
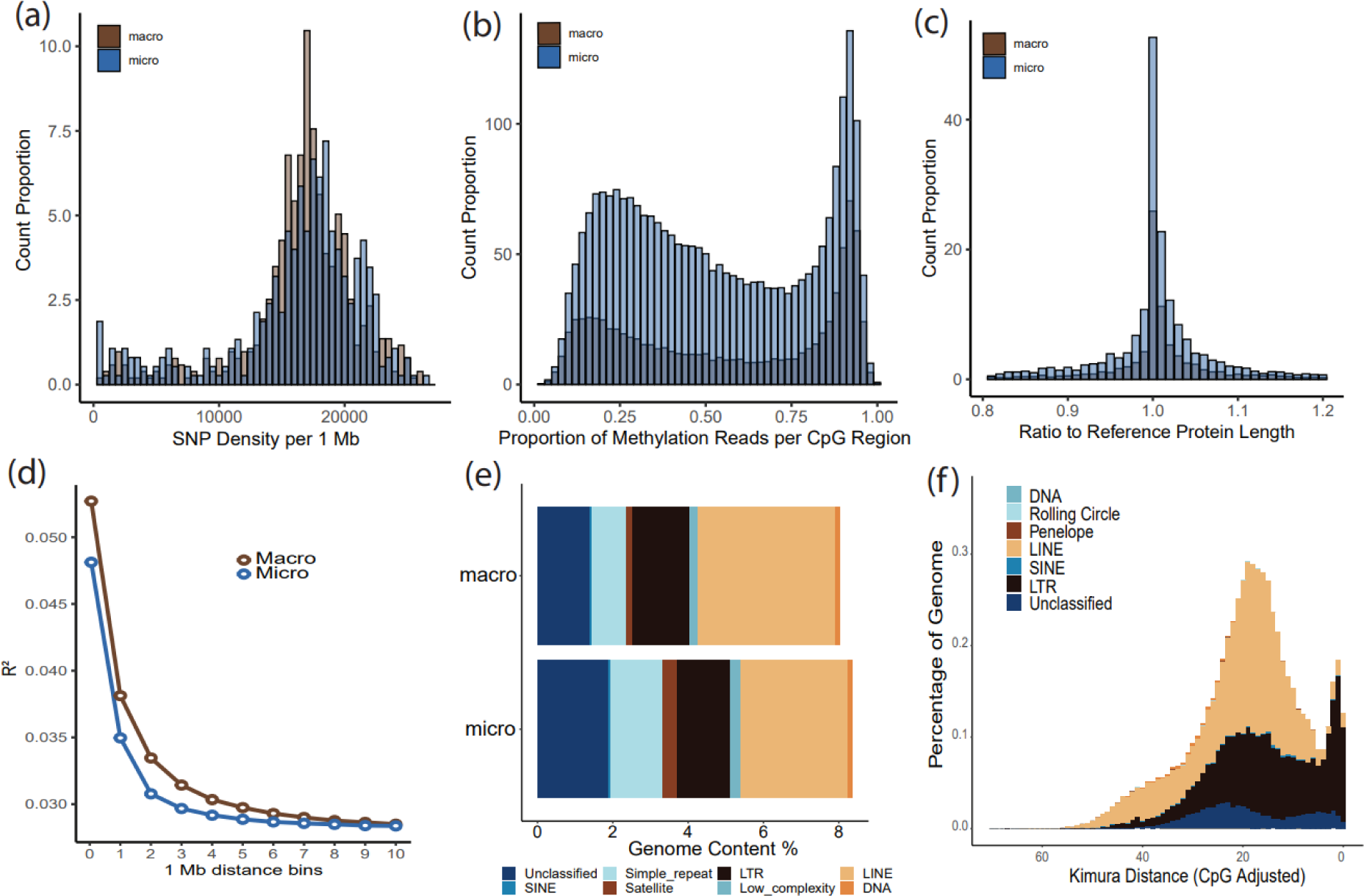
Genome summary information for the *Acridotheres tristis* AcTris_vAus2.0 genome. Panel **(a)** is the histogram of SNP density per 1 Mb window across macro and micro chromosomes. Panel **(b)** is the histogram of the proportion of methylated reads per CpG region across macro and micro chromosomes. Panel **(c)** is SAAGA annotation quality assessment of the proteome generated by the final annotation against the *Gallus gallus* reference proteome. Panel **(d)** is the average linkage disequilibrium (R^2^) in 1 Mb bins for macro and micro chromosomes. Panel **(e)** is the REPEATMASKER annotation of the macro and micro chromosomes separately using the custom assembled repeat library. Panel **(f)** is the EARLGREY plot of Kimura Distances for the different classes of TEs across the genome.

From our final annotation of 19,836 genes, we plotted gene density to reveal genomic regions of interest (Fig. 3; track 2). For instance, some gene density peaks aligned with highly non-variant regions (e.g. ∼20% into chromosome 1), which suggests the genes contained within this region are highly conserved, indicating some level of purifying selection against novel variation (Cvijović *et al*. 2018). Alternatively, some regions of high gene density coincide with regions of high variant density (e.g. start of Z chromosome), pointing to high inter-individual diversity and diversifying selection (Lavretsky *et al*. 2015). Gene density was generally higher on microchromosomes than macrochromosomes. Predicted gene sequences across these two groups of chromosomes had similar quality profiles as indicated by comparisons to the *Gallus gallus* reference proteome (Fig. 4c).

Profiling repeats across the common myna genome, we identified more repeat coverage on microchromosomes, though the difference was minimal (Fig. 4e). The reduced repeat content of the macrochromosomes and microchromosomes (Fig. 4e) compared to the overall assembly (Fig. S10c) indicates that the contig fragments not incorporated into the primary superscaffolds of the assembly are very high in repeat content, as would be expected for hard to assemble regions (Peona *et al*.

2018). We further examined TEs across the genome, finding that a majority of Tes were long interspersed nuclear elements (LINE); specifically chicken repeat 1 (CR1), and long terminal repeats (LTR); specifically endogenous retroviruses (ERV) (Fig. 4d&e, Fig. S10), reflecting similar profiles found in other avians (Gao *et al*. 2017). The largest peak of TE expansion was mostly driven by LINE/CR1 elements, coupled with LTR/ERVL (Fig. 4f). Conversely, the more recent and smaller burst of Tes was dominated by several groups of ERVs, specifically and in order of contribution: ERVK, ERV1, and ERVL (Fig. 4f). This most recent burst of Tes has been seen in other avian species (Bailey *et al*. 2023), though is unusual as most have TE peaks at greater Kimura distances (Kapusta & Suh 2017; Prost *et al*. 2019).

Using this ONT data we quantified methylation proportions at 18,501,863 CpG sites across the genome. The methylated CpG site density was generally higher on microchromosomes compared to macrochromosomes (Fig. 3; track 3), though with similar profiles (Fig. 4b). A high density of highly methylated reads (75%+ methylated) is noted in several genomic regions, such as midway through Chr 1A and 4A, and throughout the major sex (Z) chromosome (Fig. 3; track 3). These CpG sites were then summarised into 175,596 methylated regions with an average of 36 methylated sites over an average of 370 bp. Of these methylated regions, 9,524 occurred 5kb (or less) upstream of gene start sites, and showed signals of hypomethylation (35.8% of reads methylated) compared to the genome wide average of methylated reads per region (48.1%). In contrast to these gene-associated methylation patterns, we found hypermethylation in methylated regions overlapping transposable elements (63.0% of reads methylated). The common myna methylome has a similar level of overall CpG methylation, as well as gene- and TE- associate patterns, compared to other avian species, reflecting the role CpG modifications play in silencing TE transcription and regulating gene transcription (Li *et al*. 2011; Derks *et al*. 2016). Interspecific differences in methylome patterns are hard to directly compare, because these will be highly impacted by different methylation profiling methods and analytical decisions (Viitaniemi *et al*. 2019; Höglund *et al*. 2020; Bailey *et al*. 2023). However, profiling these patterns in the common myna lays important groundwork for future studies into genome-methylome interactions (e.g. Sun *et al*. 2021; Sepers *et al*. 2023).

Using LD based inference, we resolved the linear recombination profile of each autosomal chromosome. We identified that macrochromosomes had consistently higher recombination at telomeric ends, while recombination patterns along the microchromosomes were less predictable and often higher in mid-chromosomal regions (Fig 4a; track 5) reflecting trends seen in other avians (e.g. Groenen *et al*. 2009). Pairwise linkage disequilibrium decay over macrochromosomes and microchromosomes were similar, though with the former having consistently higher average R^2^ scores until background recombination values were reached at a binned distance of approximately 10 Mb (Fig. 4d). Avian genomes generally report stronger linkage disequilibrium on macrochromosomes compared to microchromosomes (Stapley *et al*. 2010; Fu *et al*. 2015), though we note that some differences in linkage disequilibrium patterns for immediately adjacent markers (Stapley *et al*. 2010) may be impacted by analytical methods and marker density filtering.

### 3.4 Gene family expansions in *Acridotheres tristis*

We used ORTHOFINDER and CAFE5 to summarise gene orthogroups over 12 species and identify expansion and contraction events. We identified a total of 18,659 gene families across the 12 species included in our analysis. Specifically, the common myna genome had a larger number of significant gene families that were lost compared to gained (Fig. 5). A similar pattern is seen in the most closely related species the European starling, and indeed appears to be the most common state of avian proteomes across the species included in our analysis (Fig. 5) and previous similar analysis (Liu *et al*. 2022; Karawita *et al*. 2023) owing to the gene loss that characterises avian genomes (Zhang *et al*. 2014). When considering all gene family changes (not just significant ones) gene family expansion and contraction was more even, a trend not seen in the European starling nor most other avian species (Fig. S12). The only avian included in our analysis that reported more gene family expansions than contractions was the zebra finch (Fig. 5 & Fig. S12). This pattern is seen in other studies of *T. guttata* (Liu *et al*. 2022; Karawita *et al*. 2023), and is not seen in the zebra finch’s nearest relative *Lonchura striata domestica* despite only 10 million years’ divergence (Kumar *et al*. 2022), though this result may also be an artifact of differing annotation approaches.

**Figure 5.**
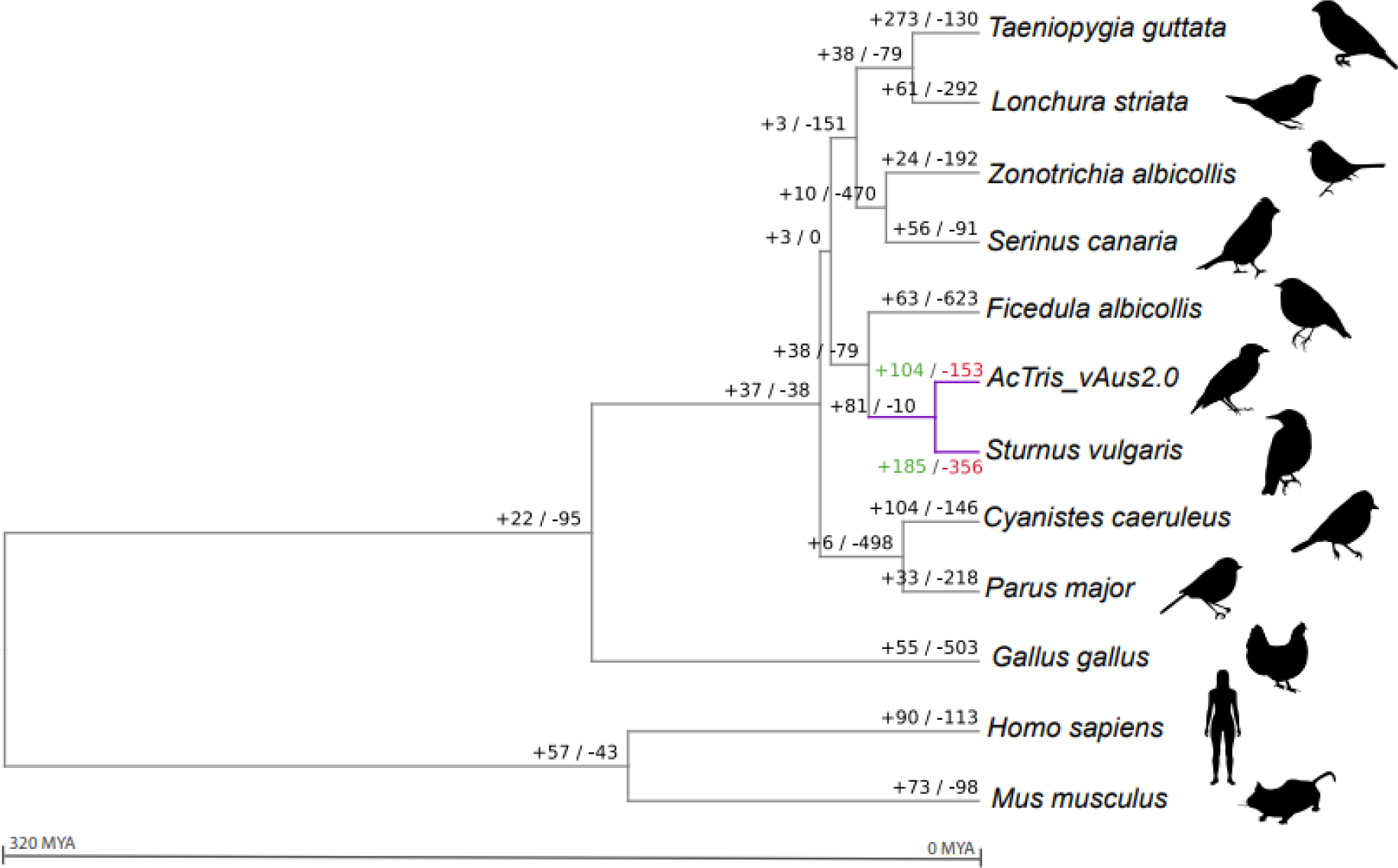
Gene family expansion analysis across Aves species, and within *Acridotheres tristis*. This figure depicts the total number of gene families (specifically, significant hierarchical orthogroups or HOGs) that had expanded or contracted within branches. The green (+) and red (−) numbers represent the expanded and contracted gene families, respectively. Sturnidae lineage indicated in purple. Pictures taken from https://www.phylopic.org/.

We then investigated the gene function for common myna gene families that had undergone significant expansion or contraction, using GO terms annotated by INTERPROSCAN. Expanded gene families were associated with biological processes such as transcription and protein modification, signalling pathways, and regulation of DNA-templated transcriptome, membrane adhesion and vesicle transport (Fig. S13). Contracted gene families were associated with methylation, viral processes, transcription, and nervous system development (Fig. S13). We note that in some cases the same gene ontology group appeared in both expanding and contracting families, because expansions and contractions are calculated based on HOGs and thus annotated GO terms are not guaranteed to be unique. Appearance of the same GO term in both expanding and contracting gene families was seen for some common biological processes and may indicate higher redundancy in those biological pathways (Nowak *et al*. 1997; Delattre & Félix 2009).

We note an interesting trend during examination of the gene families that underwent expansion and contraction across the different species and linages assessed. Many of the gene families that underwent the largest contraction in the common myna were noted to have expansions in the nearest sister taxa, the European starling (Fig. 5, Table S7). Conversely, gene families undergoing expansions in the common myna were often found to be contracting within the European starling. Both the common myna and European starling (Stuart *et al*. 2022a) genomes were annotated in similar ways using the same reference proteomes to inform the homology-based annotation tool (GEMOMA) alongside *ab-initio* gene discovery, and the longest isoform proteomes contained a similar number of gene sequences (19,836 for common myna to 21,843 for European starling) and annotation BUSCO completeness (98.5% to 98.2%). While many of these sequences were unable to be annotated by INTERPROSCAN, (indicating they are likely novel and/or lineage specific sequences), some biological processes (e.g. common myna expanding GO:0003677 - DNA binding, and common myna contracting GO:0006955 - immune response, GO:0042613 - MHC class II protein complex) were identified (Binns *et al*. 2009). These flagged gene families may act as shortlists for biologically interesting gene families that may be rapidly evolving across the Sturnidae lineage.

### 3.5 Demographic history inference

We used two different historic demographic statistical approaches, because the sequentially Markovian coalescent methods in PSMC is more useful for demographic changes in the distant past while site frequency spectrum based STAIRWAY PLOT performs better over more recent generations (Patton *et al*. 2019). Using PSMC to analyse demographic changes based on a singular genome, we observed that many of these individuals’ genomes signalled a large increase in effective population size ∼60 kya (Fig. 6a), followed by a sharp population decrease ∼10-20 kya. This population boom exists roughly after marine isotope stages MIS4 (De Deckker *et al*. 2019), and before MIS2, the last glacial maximum (Lisiecki & Raymo 2005). This pattern is similar to that in some other Eurasian passerine species over this time period (Nadachowska-Brzyska *et al*. 2015; Nadachowska-Brzyska *et al*. 2016), though we note that this result is purely correlative and highly dependent on generation time and mutation rate choices. We observe also that the size of the population increase is highly variable across analysed genomes and curiously, individuals from more bottlenecked invasive populations produced dramatically larger population expansion estimates compared to native range individuals despite the ancestors of both populations presumably occurring in the native range at the time of population expansion (Atsawawaranunt *et al*. 2023). This is highly suggestive that recent demographic shifts may introduce systemic bias in coalescent estimates of historical effective population sizes, thus signalling caution when interpreting relative signals in highly bottlenecked populations.

**Figure 6.**
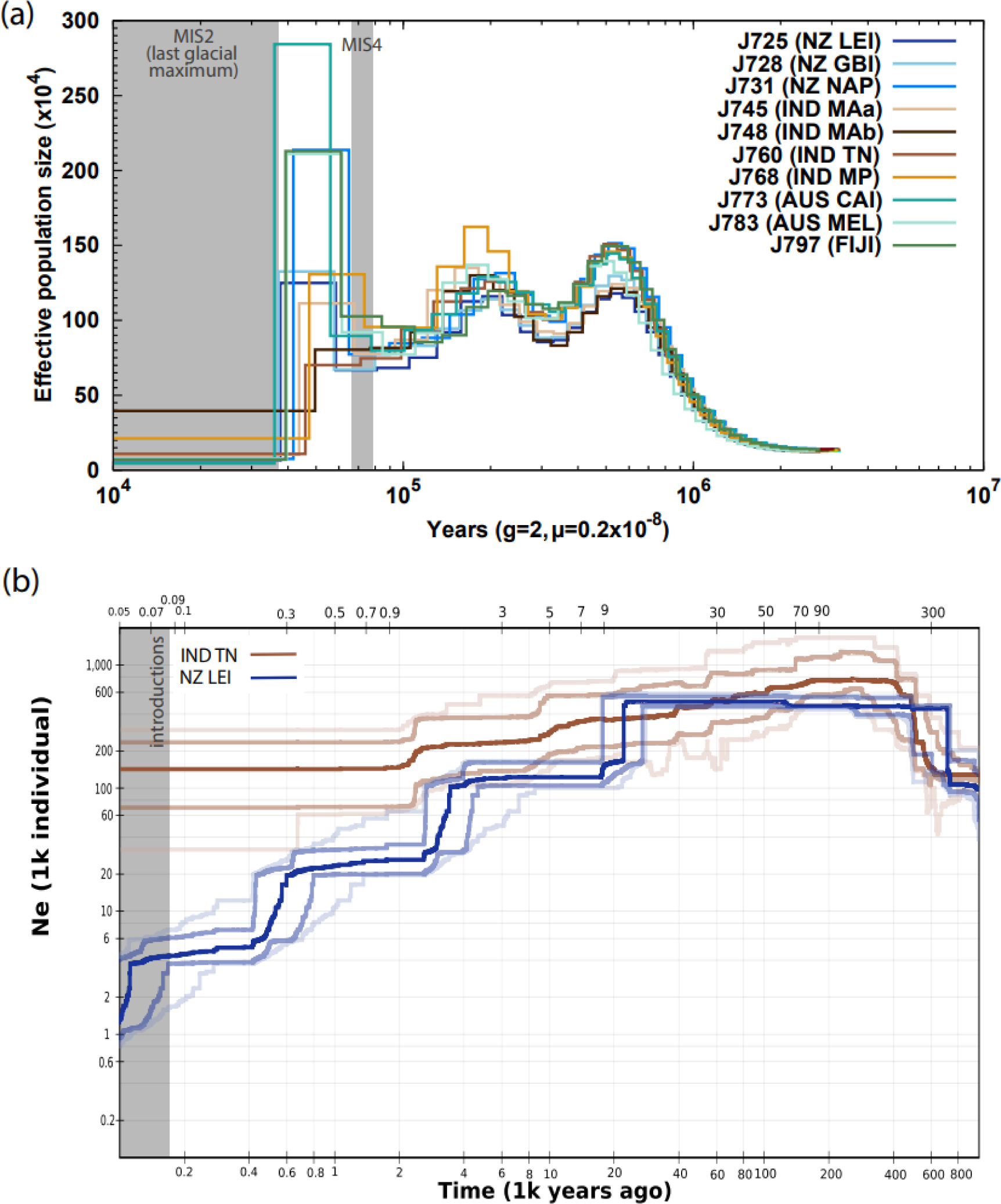
Demographic history of *Acridotheres tristis*. Panel (a) depicts the ancient demographic history as generated by PSMC from the resequenced whole genomes of 10 individuals, with native range individuals (IND) in browns, and invasive range individuals (FIJI, AUS, and NZ) in blue and green. Panel (b) depicts the more recent demography as generated by STAIRWAY PLOT from individuals from one native range sample site and one introduced range sample site (IND TN N = 8, NZ LEI N = 8; lighter lines indicate 75% and 95% confidence intervals). Full sample site names are given in Table S3.

This paradox might alternatively be explained through the observation that this dramatic population expansion was not ubiquitous across all samples analysed, and indeed was not present in some of the native range individuals. The mixed signal for population expansion within the native range was evident particularly when bootstrapping was performed (Fig. S14). When the number of native range samples analysed were increased (Fig. S15), we noted that the mixed population expansion signal was visible in all native range sample sites except for Maharashtra subpopulation A (MAa), a sample site previously observed to be more bottlenecked than the rest of the native range and which clustered closely with some invasive populations (Atsawawaranunt *et al*. 2023) (though we acknowledge small sample size in this analysis limits the strength of this conclusion). This result is indicative of two things. Firstly, that this population expansion event may not have covered the entire native range but possibly differed based on climate or biome differences throughout Eurasia across MIS4-MIS2 (Ray & Adams 2001; Guo *et al*. 2019). Secondly, the lack of a mixed signal in the invasive range samples analysed (Fig. 5a) supports previous conclusions that it is likely that the ancestors of invasive populations were sourced from similar locations for each separate introduction effort (Atsawawaranunt *et al*. 2023).

When demographic patterns over more recent generations were estimated using the population-based site frequency spectrum approach employed by STAIRWAY PLOT, both the native and invasive range population produced an estimated peak effective population size of 400,000-600,000 individuals, followed by a slow decline towards present day (Fig. 6a). While both populations reported an increase in population size at around 600 kya, this decreased within the first 10 steps of both population’s plots, and it is best practice to not overinterpret this region (Liu & Fu 2015). This increase does however correspond to a similar time frame of a population increase reported by PSMC. In both analyses, we observe that the bottlenecked invasive populations had a more rapid estimated rate of demographic change. Despite these populations only having diverged within the last 200 years (Beesley *et al*. 2023), the invasive NZ LEI population results in a more pronounced effective population size decrease towards present day time. Interestingly, the initial point of effective population size decline estimated by this population corresponds to the PSMC estimated decline at 20 kya. However, NZ LEI continues to decline sharply until present day and produces more recent population size estimates of 2,000 individuals, which is well below that estimated by IND TN (160,000) or both native and introduced individuals in PSMC (50,000-400,000).

## 4 CONCLUSION

Here we present a chromosome level assembly of the common myna, *A. tristis*, one of the most successful invasive avian species recorded over the past 200 years. There are substantial resources and efforts going towards understanding this species in the regions it has become invasive and we believe genomic insights will provide valuable tools towards those efforts. Conversely, as increasing numbers of species suffer reduced population sizes, successful introduced/invasive species like the common myna provide unique insight as to how some species can successfully inhabit a diversity of ecological conditions, in this case from a small founder population size. This high-quality chromosome level assembly and annotation will provide valuable genomic context for future studies on this species. Through this work we describe the genomic landscape of this species, including genome wide allelic diversity, methylation, repeats, and recombination, as well as an examination of gene family expansions and contractions. Using demographic analysis we provide the first whole genome level insight into this species through ancient and recent time. We identify that some native regions underwent a dramatic population increase between the two most recent periods of glaciation, but also reveal artefactual impacts of bottlenecking on demographic analysis.

## Data Accessibility

All raw data will be available via NCBI. Code is available at GitHub (https://github.com/katarinastuart/At1_MynaGenome).

## Acknowledgments

We extend many thanks to Dinindu Senanayake and Joseph Guhlin for their technical support for analyses. Huge thanks to Andrew King for sample preparation and extraction. We are very grateful to Aaron Darling, Dominique Gorse and their groups for their work in sequencing and assembling an earlier version of the myna genome. Thanks to Stella Loke for hosting A.W. and K.C.S. for ONT training. And our thanks to Australia Museum staff including Leah Tsang, Scott Ginn and Tracey McVea for their support in sample loans and research collaboration contracts. Finally, our thanks to Mark Peck at the Royal Ontario Museum and to the many individual collectors in Australia and New Zealand who contributed myna samples.

We gratefully acknowledge funding from The New Zealand Royal Society Te Apārangi Marsden Grant (grant number UOA1911), Australian Museum Foundation and City of Sydney Environmental Grants Program. Seed funding was also generously provided by the University of Auckland’s Faculty of Science Sustainability Theme, Computational Biology Theme and the Digital Biology Interface Institute. A University of Auckland Doctoral Scholarship supports K.A.

## Author contributions

A.W., K.C.S., R.N.J., R.M., L.A.R. and A.W.S. designed the research, A.W. conducted the ONT sequencing and led the myna genome assembly, K.C.S. led the genome analyses. R.M. and K.M.E. coordinated data collection from Australia for the genome individual and population samples. A.W.S. coordinated data collection for New Zealand with K.A. contributing insights on the global population structure. K.C.S. led the writing of the paper, with input from A.W.S. and feedback from all authors. All authors read and approved the final manuscript.

