## Supplementary Material for "The genome of a globally invasive passerine, the common myna (*Acridotheres tristis*)"

**Supplementary materials**

### Appendix 1: Sampling, extraction, and sequencing

*Extractions*

High-molecular weight gDNA was extracted using the QIAGEN PureGene Tissue Extraction protocol for linked-read sequencing, while for ONT sequencing, gDNA was extracted with the Qiagen Genomics Tips and buffer set for the first round of sequencing. Subsequent iterations of ONT sequencing used genomic DNA extracted with the PacBio Nanobind BigDNA animal tissue kit and further purified using the Short Read Eliminator reagent after extraction. Messenger RNA was extracted with the Qiagen RNeasy Plus Universal Mini Kit and Qiagen Oligotex mRNA Mini Kit. Nucleic acid concentration and quality was assessed using a NanoDrop, Qubit and TapeStation.

*gDNA and cDNA sequencing*

High molecular weight gDNA (1 ug) was prepared for 10x Chromium linked-read sequencing according to the manufacturer's recommended protocols. A 10x GEM library was barcoded using the Chromium Genome Reagent Kits (v2 Chemistry). The library was run on a single lane of a S4 flowcell and sequenced using the Illumina Hiseq X Ten sequencing platform (150 bp paired end reads) at Macrogen, Seoul, Republic of Korea.

Oxford Nanopore Technologies ligation sequencing libraries were prepared with 1.5ug gDNA per library. An initial round of sequencing using the LSK109 library kit, subsequent rounds used the LSK110 and LSK112 releases (Table S2). All sequencing was performed on R9.4.1 flow cells. A total of 6 flow cells were used. Libraries were loaded (at ~50fmol) multiple times per flow cell, with nuclease flushes to restore pore functionality between loads.

The cDNA was quantified using an Agilent 2200 TapeStation (Agilent, California, USA) and Nanodrop spectrophotometer (Thermo Fisher Scientific). cDNA from three tissues (liver, heart, testes) was then prepared for sequencing using a TruSeq stranded mRNA library prep kit, and sequenced on a NextSeq 75bp PE high output sequencing run at the Ramaciotti Centre for Genomics (UNSW, Sydney, NSW).

**Appendix 2: Manual genome curation**

The initial nanopore assembled and polished genome assembly was mapped, using MINIMAP2 v2.24 (Li 2018), to the zebra finch (*Taeniopygia guttata)* genome assembly (GCF_003957565.2), as well as a Vertebrate Genomes Project (VGP) common myna genome assembly (GCA_027559615.1). These alignments were visualised as dot plots using DGENIES v1.4.0 (Cabanettes & Klopp 2018), and the dot plots manually inspected for signatures of misassemblies (Fig. S1). To investigate these putative misassemblies, we used MINIMAP2 to map reads (raw ONT, and trimmed 10x reads) to the assembly so coverage around potential misassembly points could be investigated. In addition to the raw reads, we aligned the assemblies of the European starling (*Sturnus vulgaris*) genome (GCA_023376015.1) (Stuart *et al.* 2022a) to provide extra context around potential misassembly points. These three tracks were visualised in IGV v2.15.4 (Robinson *et al.* 2011) and locations of misassembly signatures were examined. Contigs that contained evidence for potential misassembly (for examples, see: Fig. S2) were manually broken using BEDTOOLS V2.30 *getfasta* (Quinlan & Hall 2010). We used DIPLOIDOCUS v1.1.1 (Chen *et al.* 2022) to check for assembly artefacts (runmode=purgehaplotig) and vector contamination (runmode=vecscreen using the NIH UniVec database). NUMTFINDER v0.5.1 (Edwards *et al.* 2021) was used to identify the mitochondrial genome and search for misassemblies of it, using the common myna mitochondrial genome as a reference (CM050619.1). Contigs below length of 1,000bp were identified using BBMAP v38.81 (Bushnell 2014) and removed. BLOBTOOLS v1.0 (Laetsch & Blaxter 2017) was used to assess if any of the assembled contigs may have belonged to contaminants in the raw sequencing data, using BLASTN v2.13.0 (Camacho *et al.* 2009) (-task megablast) to assemble the database, and the ONT reads mapped to the genome using MINIMAP2. Sequences that were identified as contaminants were removed from the assembly (Fig. S3).

**Appendix 3: Z chromosome genomic rearrangement**

Prior to use as a reference for synteny scaffolding, the VGP *A. tristis* genome was examined for synteny against the *T. guttata* genome using MINIMAP2 and visualised in DGENIES (Supplementary materials: Fig. S5). The dot plot revealed that approximately 17 Mb section of a *T. guttata* autosomal chromosome was assembled onto the end of the Z chromosome of VGP *T. guttata*. To investigate whether this was a genuine genomic rearrangement in *A. tristis* relative to the zebra finch or a misassembly, we mapped our raw ONT reads to VGP *T. guttata* using MINIMAP2 and visualised in IGV (Supplementary materials: Fig. S5). High ONT read concatenation around the scaffold join site indicated this was likely a misassembly in the VGP assembly. We took a conservative approach and excluded the equivalent region in our assembly from synteny based scaffolding, but during scaffold renaming renamed it based on *T. guttata* synteny.

**Table S1 | *Acridotheres tristis* individuals whole genome resequenced.** SVC = single nucleotide variant calling, RC = recombination profiling, DP = demography PSMC, DS = demography stairway plot.

| Country & Status | Population | Abbreviated ID (internal) | Full name (internal) | SRA Accession | Analysis used |
|---|---|---|---|---|---|
| INDIA *Native* | Tamil Nadu (TN) | J753 | IND_TN_3695_M0298 | *forthcoming* | SVC, RC, DS |
| | | J754 | IND_TN_3696_M0299 | *forthcoming* | SVC, RC, DS |
| | | J755 | IND_TN_3697_M0300 | *forthcoming* | SVC, RC, DS |
| | | J756 | IND_TN_3699_M0302 | *forthcoming* | SVC, RC, DS |
| | | J757 | IND_TN_3700_M0303 | *forthcoming* | SVC, RC, DS |
| | | J758 | IND_TN_3701_M0304 | *forthcoming* | SVC, RC, DS |
| | | J759 | IND_TN_3710_M0305 | *forthcoming* | SVC, RC, DS |
| | | J760 | IND_TN_3711_M0306 | *forthcoming* | SVC, RC, DS, DP |
| | Madhya Pradesh (MP) | J761 | IND_MP_3751_M0317 | *forthcoming* | SVC, RC |
| | | J762 | IND_MP_3752_M0318 | *forthcoming* | SVC, RC |
| | | J763 | IND_MP_3753_M0319 | *forthcoming* | SVC, RC |
| | | J764 | IND_MP_3754_M0320 | *forthcoming* | SVC, RC |
| | | J766 | IND_MP_3756_M0322 | *forthcoming* | SVC, RC |
| | | J767 | IND_MP_3757_M0323 | *forthcoming* | SVC, RC |
| | | J768 | IND_MP_3758_M0324 | *forthcoming* | SVC, RC, DP |
| | Maharashtra subpop A (MAa) | J745 | IND_MA_979_M0213 | *forthcoming* | DP |
| | Maharashtra subpop b (MAb) | J748 | IND_MA_3731_M0307 | *forthcoming* | DP |
| Fiji *Invasive* | Fiji (FIJI) | J797 | FIJI_923_M0203 | *forthcoming* | DP |
| Australia (AUS) *Invasive* | Melbourne (MEL) | J783 | AUS_MEL_12408 | *forthcoming* | DP |
| | Cairns (CAI) | J773 | AUS_CAI_11606 | *forthcoming* | DP |
| New Zealand (NZ) *Invasive* | Leigh (LEI) | J718 | NZ_LEI-06_M0006 | *forthcoming* | DS |
| | | J719 | NZ_LEI-07_M0007 | *forthcoming* | DS |
| | | J720 | NZ_LEI-08_M0008 | *forthcoming* | DS |
| | | J721 | NZ_LEI-09_M0009 | *forthcoming* | DS |
| | | J722 | NZ_LEI-10_M0010 | *forthcoming* | DS |
| | | J723 | NZ_LEI-13_M0013 | *forthcoming* | DS |
| | | J724 | NZ_LEI-18_M0018 | *forthcoming* | DS |
| | | J725 | NZ_LEI-19_M0019 | *forthcoming* | DS, DP |
| | Napier (NAP) | J731 | NZ_NAP-08_M0158 | *forthcoming* | DP |
| | Great Barrier Island (GBI) | J728 | NZ_GBI-13_M0133 | *forthcoming* | DP |

**Table S2 | ONT read batches and corresponding program version numbers** used for raw data processing used at ONT assembly steps.

| LSK Chemistry | Guppy nucleotide basecalling config | Methylation basecalling config | Total length of raw reads (Mb) | Total length of filtered reads (Mb) | Average read length (Kb) |
|---|---|---|---|---|---|
| Ligation (SQK-LSK109) | dna_r9.4.1_450bps_sup.cfg | dna_r9.4.1_450bps_modbases_5mc_cg_sup.cfg | 1,459.18 | 1,211.15 | 3.11 |
| Ligation (SQK-LSK110) | dna_r9.4.1_450bps_sup.cfg | dna_r9.4.1_450bps_modbases_5mc_cg_sup.cfg | 4,531.31 | 3,561.85 | 11.22 |
| Ligation (SQK-LSK112) | dna_r9.4.1_e8.1_sup.cfg | dna_r9.4.1_e8.1_modbases_5mc_cg_sup.cfg | 22,645.27 | 18,779.21 | 7.17 |

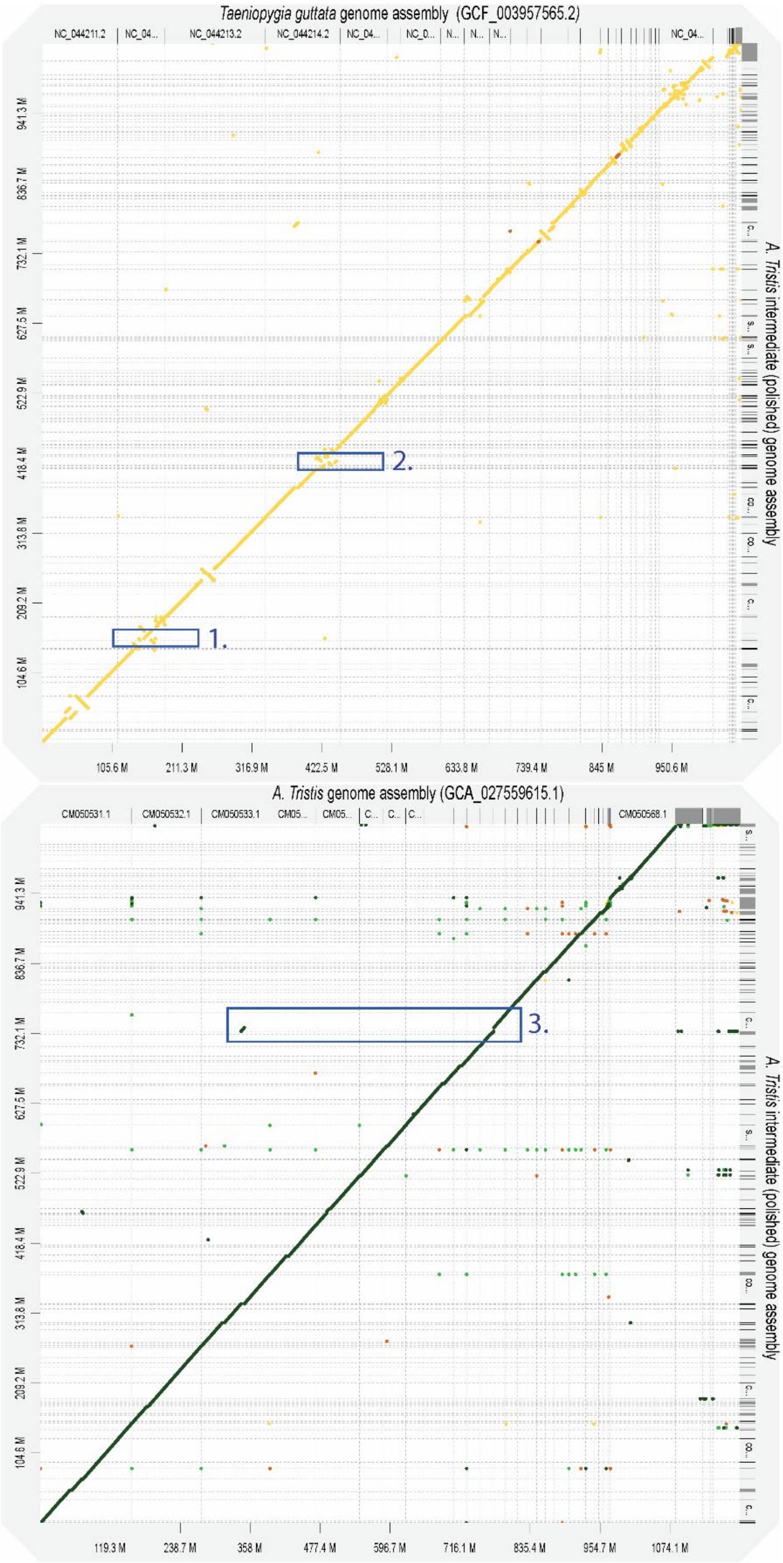


**Figure S1 | Dot plot of the polished *Acridotheres tristis* genome (initial genome assembly in this stidy) against the *Taeniopygia guttata* genome (GCF_003957565.2) and VGP *Acridotheres tristis* genome (GCA_027559615.1)**. Alignments done using MINIMAP2 v2.24 and visualised in DGENIES v1.4.0. Indicated on the dot plot are three exemplar windows that were investigated during manual genome curation that were deemed to be 1) a misassembly, 2) not a misassembly, and 3) a misassembly.

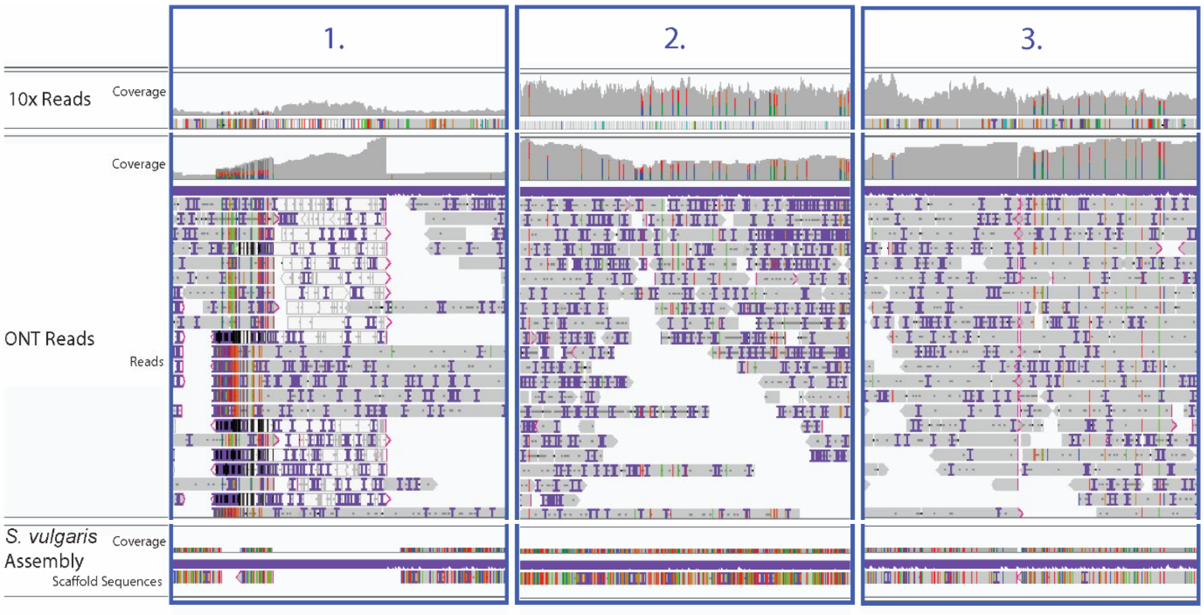


**Figure S2 | Examples of manual assembly curation**. Alignments produced using MINIMAP2 v2.24 and processed in SAMTOOLS v1.13, and finally visualised in IGV v2.15.4. Window numbers correspond to the exemplar windows from Figure S1 that were investigated during manual genome curation that were deemed to be 1) a misassembly, 2) not a misassembly, and 3) a misassembly.

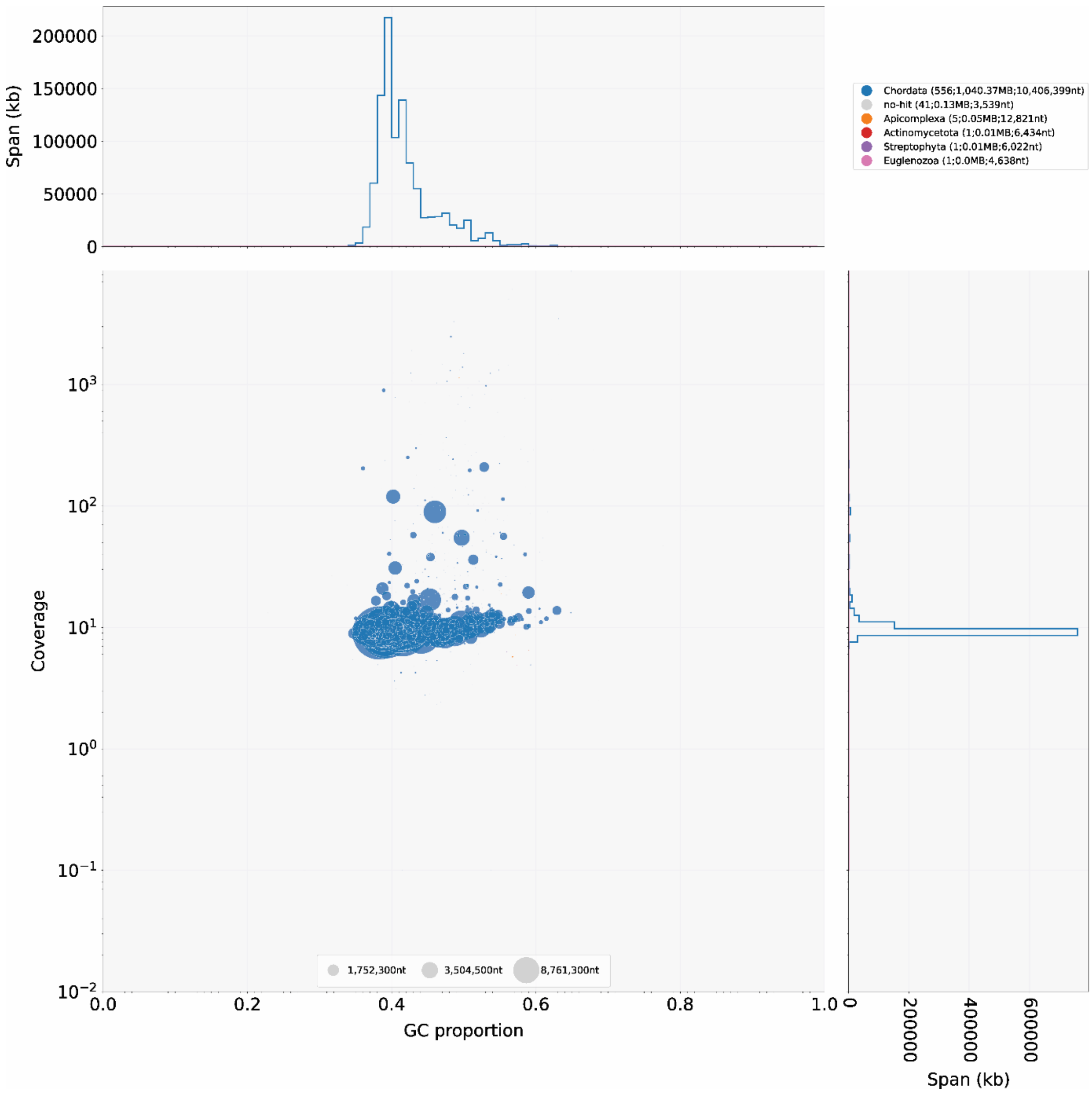


**Figure S3 | Blobtools summary plot of genome assembly contamination** produced using blast (-task megablast) to assign taxonomic identities to the sequences.

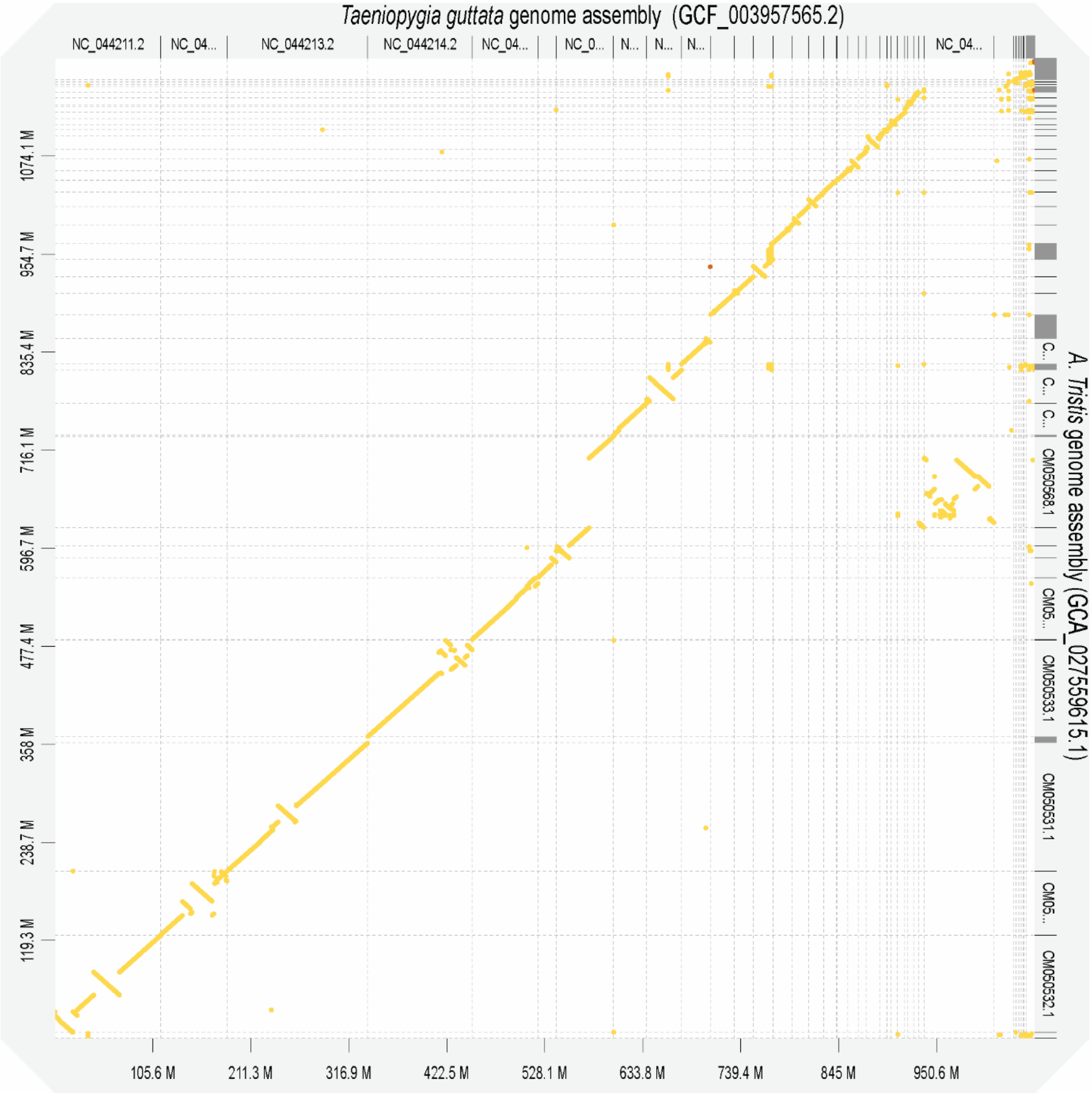


**Figure S4 | Dot plot of the VGP *Acridotheres tristis* genome (GCA_027559615.1) and *Taeniopygia guttata* genome (GCF_003957565.2)**. Alignments done using MINIMAP2 v2.24 and visualised in DGENIES v1.4.0.

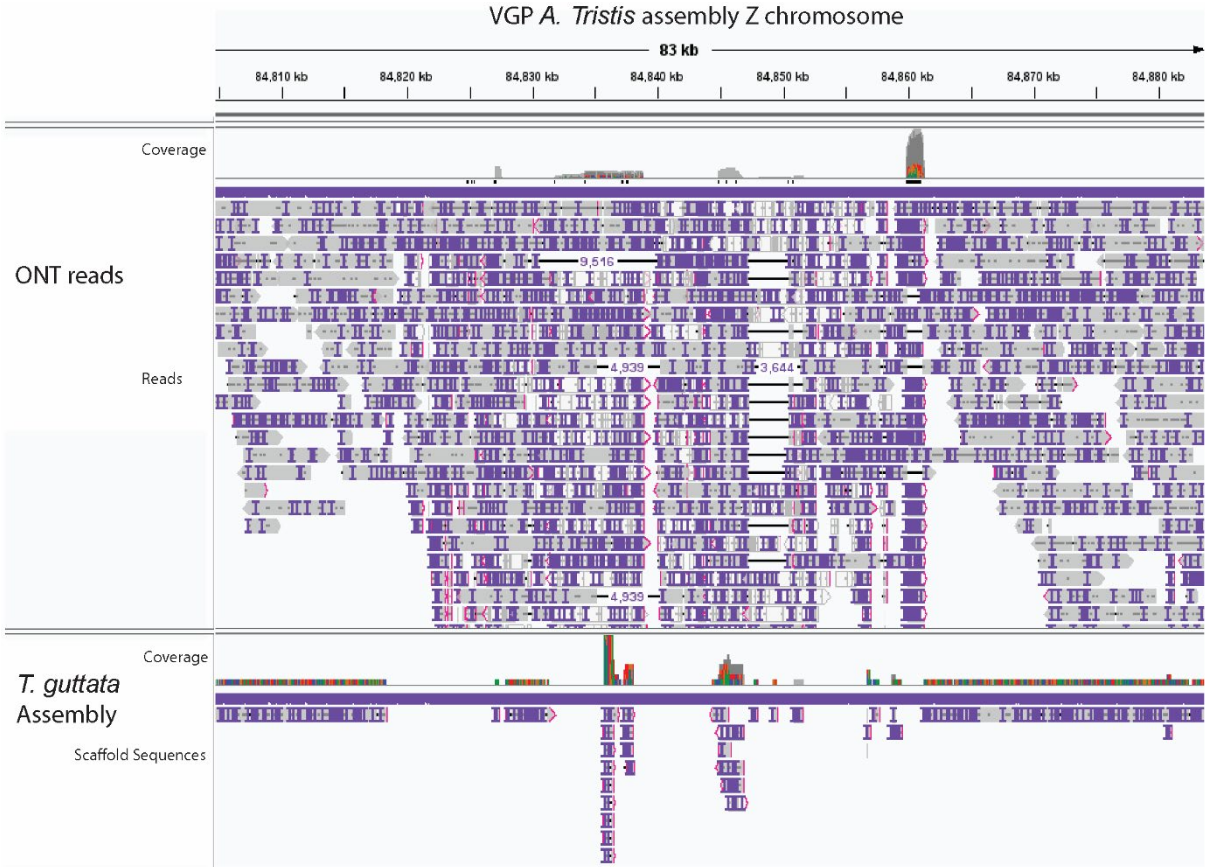


**Figure S5 | Manual examination of VGP *Acridotheres tristis* genome (GCA_027559615.1) z chromosome sequence.** Alignments produced using MINIMAP2 v2.24 and processed in SAMTOOLS v1.13, and finally visualised in IGV v2.15.4.

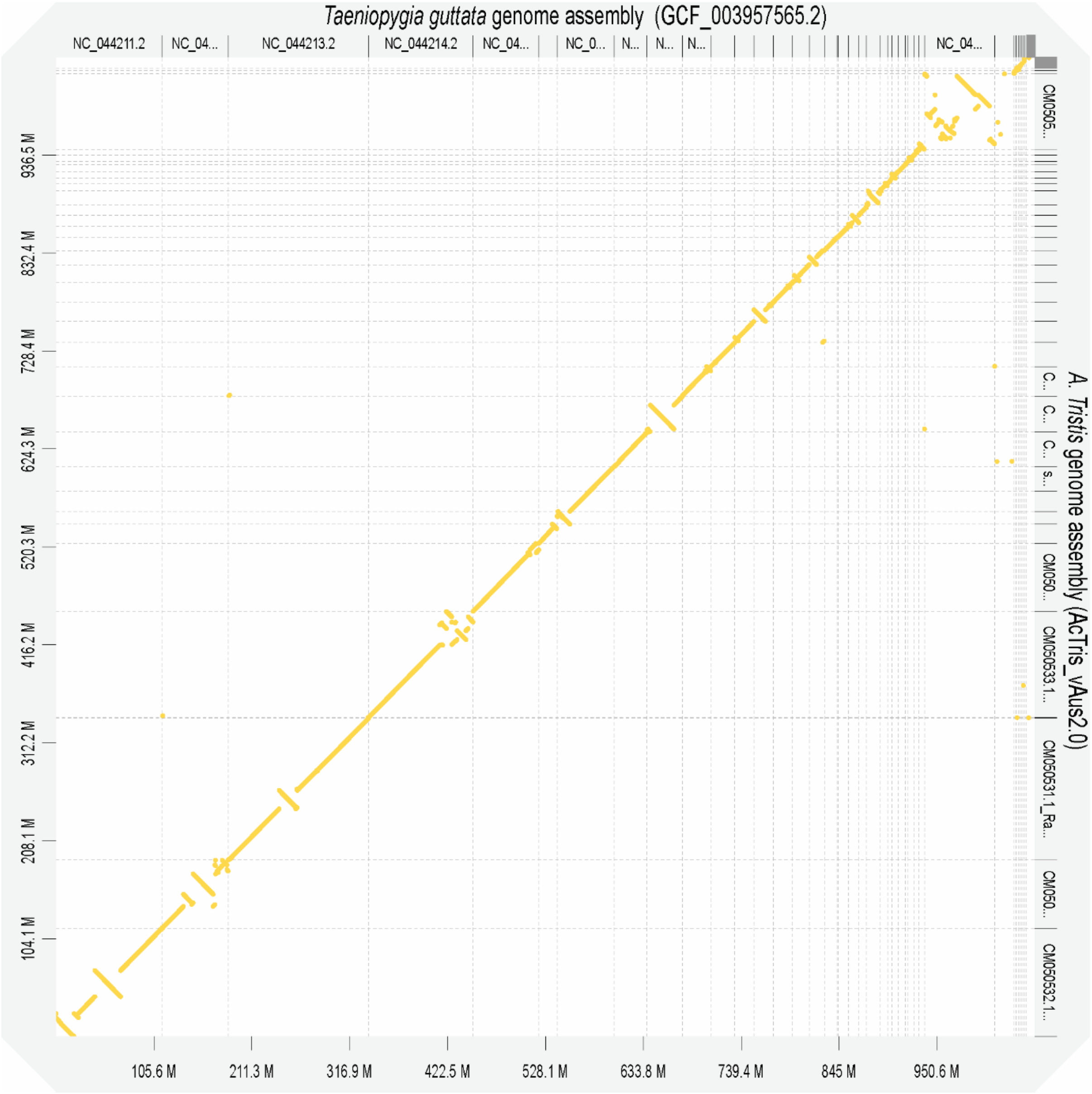


**Figure S6 | Dot plot of the *A. trisits* genome assembly (AcTris_vAus2.0) and *Taeniopygia guttata* genome (GCF_003957565.2)**. Alignments done using MINIMAP2 v2.24 and visualised in DGENIES v1.4.0.

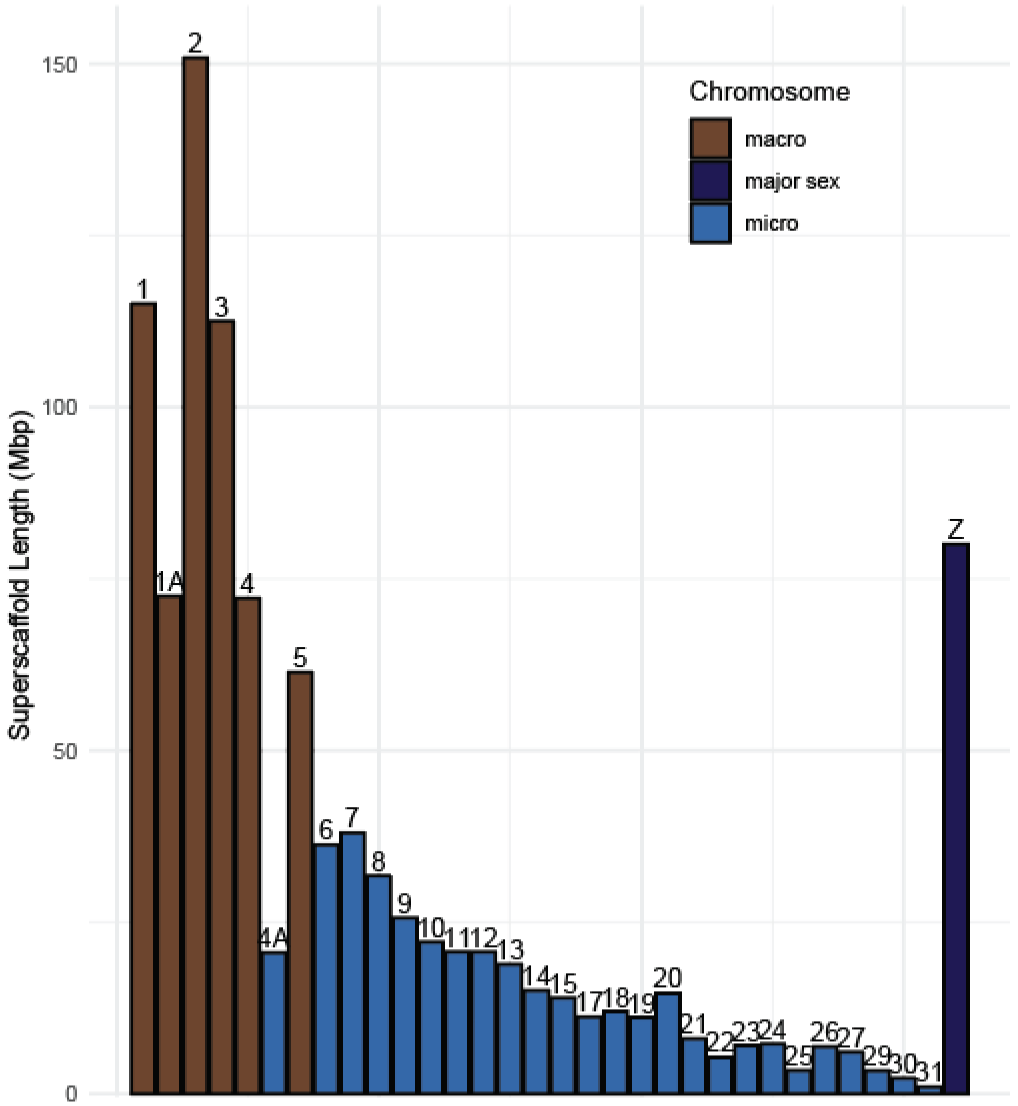


**Figure S7 | super scaffold sizes** with macro and micro chromosome numbers, based on zebra finch synteny, labelled.

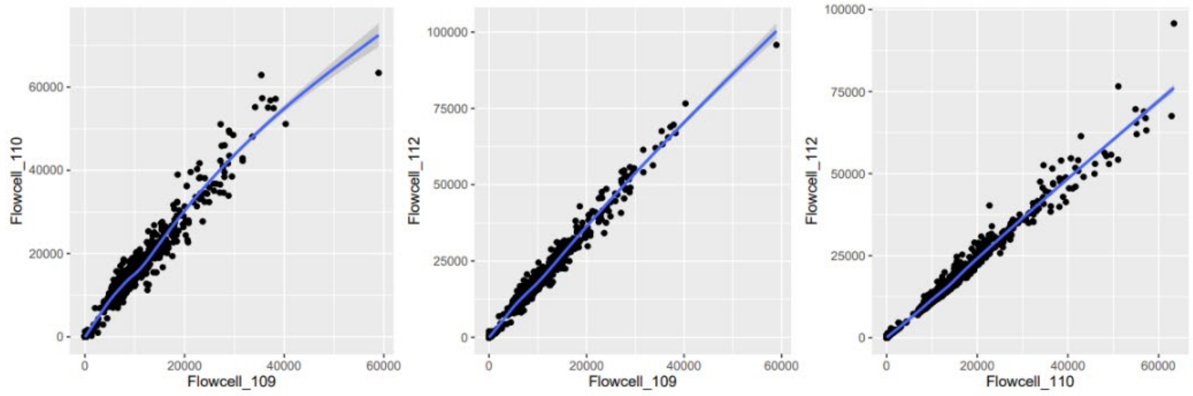


**Figure S8 | Correlation plots of the methylation calls** across the three flow cells obtained by linear regression in R.

**Table S3 | Ensembl reference species** used in the GEMOMA genome annotation.

| Common Name | Scientifiec Name | Ensembl Assembly |
|---|---|---|
| Eurasian sparrowhawk | *Accipiter nisus* | Accipiter_nisus_ver1.0 |
| Yellow-billed parrot | *Amazon collaria* | ASM394721v1 |
| Mallard | *Anas platyrhynchos* | ASM874695v1 |
| Pink-footed goose | *Anser brachyrhynchus* | ASM259213v1 |
| Swan goose | *Anser cygnoides* | GooseV1.0 |
| Greater spotted kiwi | *Apteryx haastii* | aptHaa1 |
| Little spotted kiwi | *Apteryx owenii* | aptOwe1 |
| Okarito brown kiwi | *Apteryx rowi* | aptRow1 |
| Golden eagle | *Aquila chrysaetos chrysaetos* | bAquChr1.2 |
| Burrowing owl | *Athene cunicularia* | athCun1 |
| Small tree finch | *Camarhynchus parvulus* | STF_HiC |
| Golden pheasant | *Chrysolophus pictus* | Chrysolophus_pictus_GenomeV1.0 |
| Blue tit | *Cyanistes caeruleus* | cyaCae2 |
| Emu | *Dromaius novaehollandiae* | droNov1 |
| Gouldian finch | *Erythrura gouldiae* | GouldianFinch |
| Flycatcher | *Ficedula albicollis* | FicAlb_1.4 |
| Chicken | *Gallus gallus* | GRCg6a |
| Median ground-finch | *Geospiza fortis* | GeoFor_1.0 |
| Bengalese finch | *Lonchura striata domestica* | LonStrDom1 |
| Turkey | *Meleagris gallopavo* | Turkey_2.01 |
| Great tit | *Parus major* | Parus_major1.1 |
| Ring-necked pheasant | *Phasianus colchicus* | ASM414374v1 |
| Common canary | *Serinus canaria* | SCA1 |
| African ostrich | *Struthio camelus australis* | ASM69896v1 |
| Zebra finch | *Taeniopygia guttata* | bTaeGut1_v1.p |
| White-throated sparrow | *Zonotrichia albicollis* | Zonotrichia_albicollis-1.0.1 |

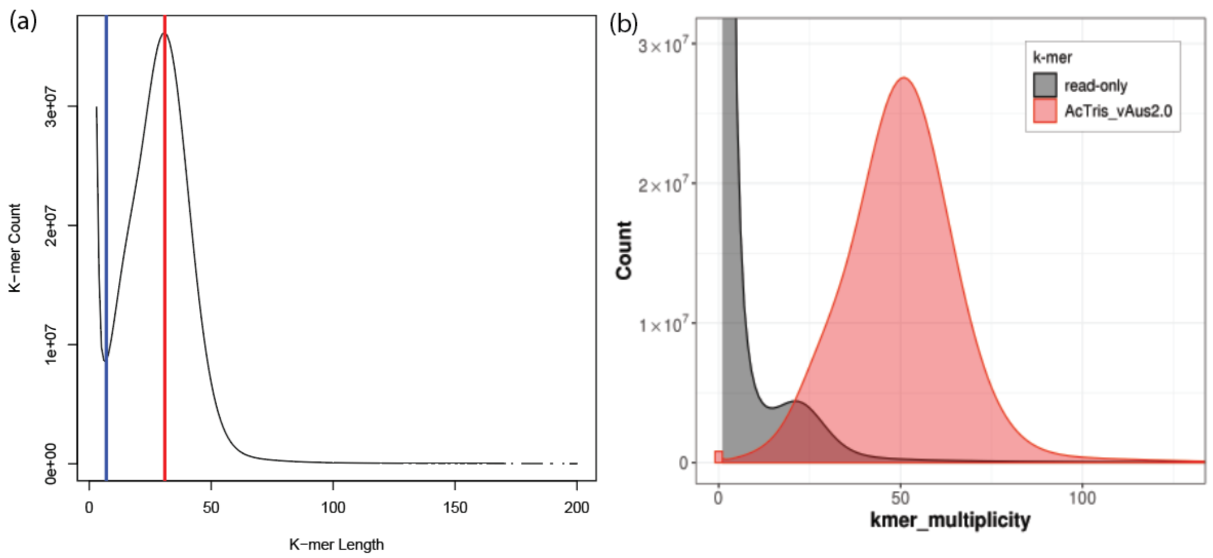


**Figure S9 | Histogram of *Acridotheres tristis* linked read gDNA k-mer counts.** Panel (a) was estimated using JELLYFISH from k=7 to k=200. Vertical red line denotes histogram peak at k=31. Genome Length = 36,028,219,169 / 31 = 1,162,200,618 bp. The final genome assembly size was used to find the using the genome size k-mer estimate. 1,040,603,622 / 1,162,200,618 = 89.54% complete. Panel (b) was estimated by MERQURY and reported a genome completeness score of 91.17% and a consensus quality value (QV) of 44.1.

**Table S4 | Genome size and completion statistics produced by JELLYFISH** for a range of initial k-mer values. Completion statistic based on the final AcTris_vAus2.0 assembly size of 1,040,603,622 bp.

| **K-mer Value** | **18** | **19** | **20** | **21** | **22** |
|---|---|---|---|---|---|
| **Predicted Genome Size** | 1,142,465,245 | 1,150,752,569 | 1,162,200,618 | 1,177,818,515 | 1,300,046,659 |
| **Completion %** | 91.08 | 90.43 | 89.54 | 88.35 | 80.04 |

**Table S5 | Genome assembly statistics for the VGP *Acridotheres tristis* genome assembly (GCA_027559615.1) for comparison.**

| Assembly Statistic | VGP *Acridotheres tristis* |
|---|---|
| Total length (bp) | 1,193,419,045 |
| Number of scaffolds | 455 |
| Scaffold N50 (bp) | 75,254,055 |
| Scaffold L50 | 6 |
| Largest scaffold (bp) | 156,254,668 |
| Mean scaffold length (bp) | 2,622,899.00 |
| Median scaffold length (bp) | 123,340 |
| Number of Contigs | 890 |
| Contig N50 (bp) | 7,308,823 |
| Contig L50 | 44 |
| Gap (N) length (bp) | 2,422,523 |
| GC content (%) | 42.32% |
| BUSCO (genome: Aves) | 8338 |
| Complete | 8094 |
| Complete (single copy) | 8060 |
| Complete (duplicated) | 34 |
| Fragmented | 46 |
| Missing | 198 |

**Table S6 | BUSCO statistics for the *Acridotheres tristis* AcTris_vAus2.0 genome annotations using BRAKER3 and GEMOMA.**

| Assembly Statistic | AcTris_vAus2.0 |
|---|---|
| BRAKER3 | |
| BUSCO (transcriptome: Aves) | 8338 |
| Complete | 7341 |
| Complete (single copy) | 5599 |
| Complete (duplicated) | 1742 |
| Fragmented | 64 |
| Missing | 933 |
| GEMOMA* | |
| BUSCO (transcriptome: Aves) | 8338 |
| Complete | 8150 |
| Complete (single copy) | 8122 |
| Complete (duplicated) | 28 |
| Fragmented | 54 |
| Missing | 134 |

* Filtered for longest transcript

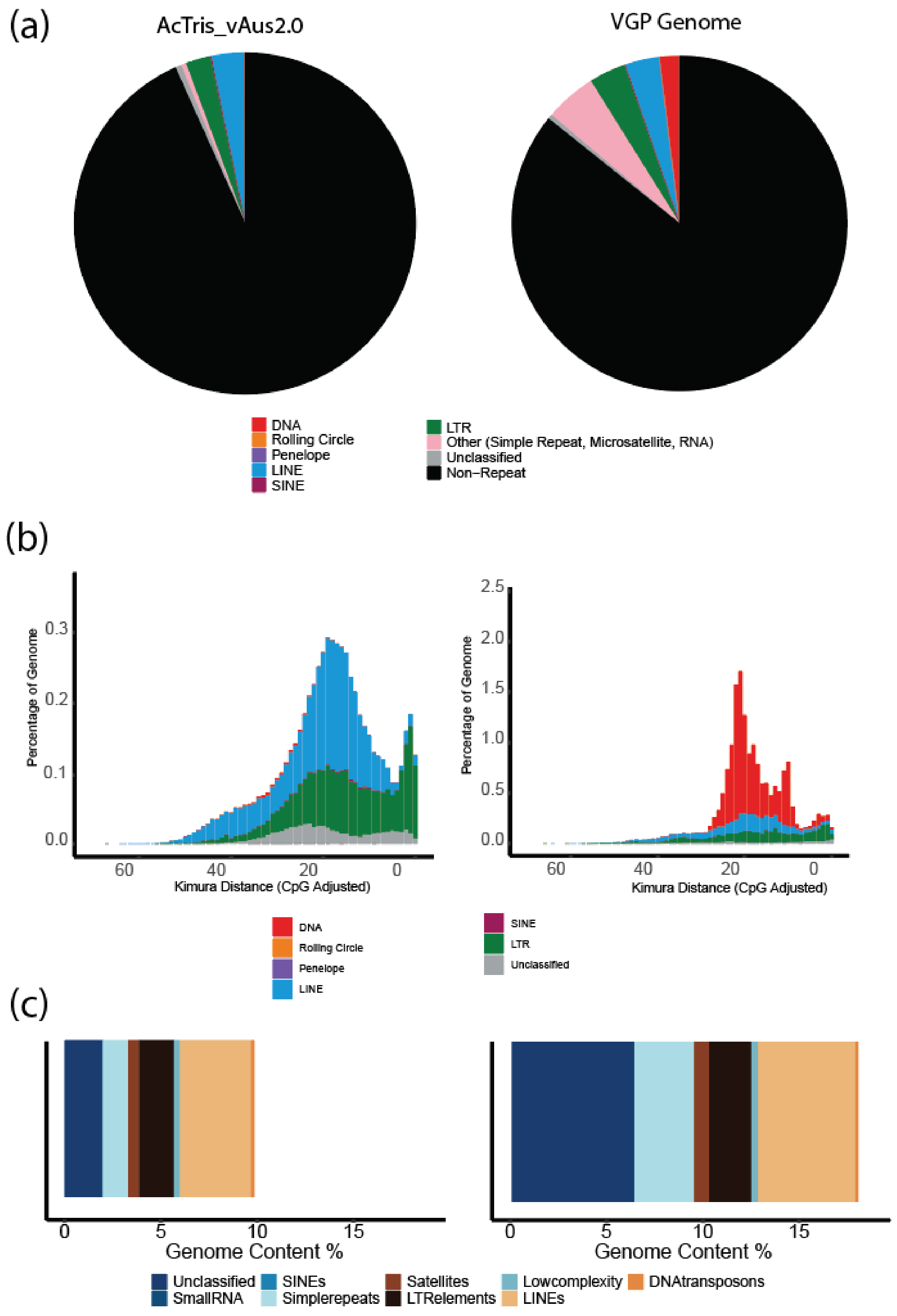


**Figure S10 | Repeat content of the *Acridotheres tristis* AcTris_vAus2.0 genome (left) and the VGP A. tristis genome (right).** Panels (a) and (b) are profiles of genome transposable element content as generated by EARLYGREY. Panel (c) is a REPEATMASKER repeat annotation of the two genome assemblies.

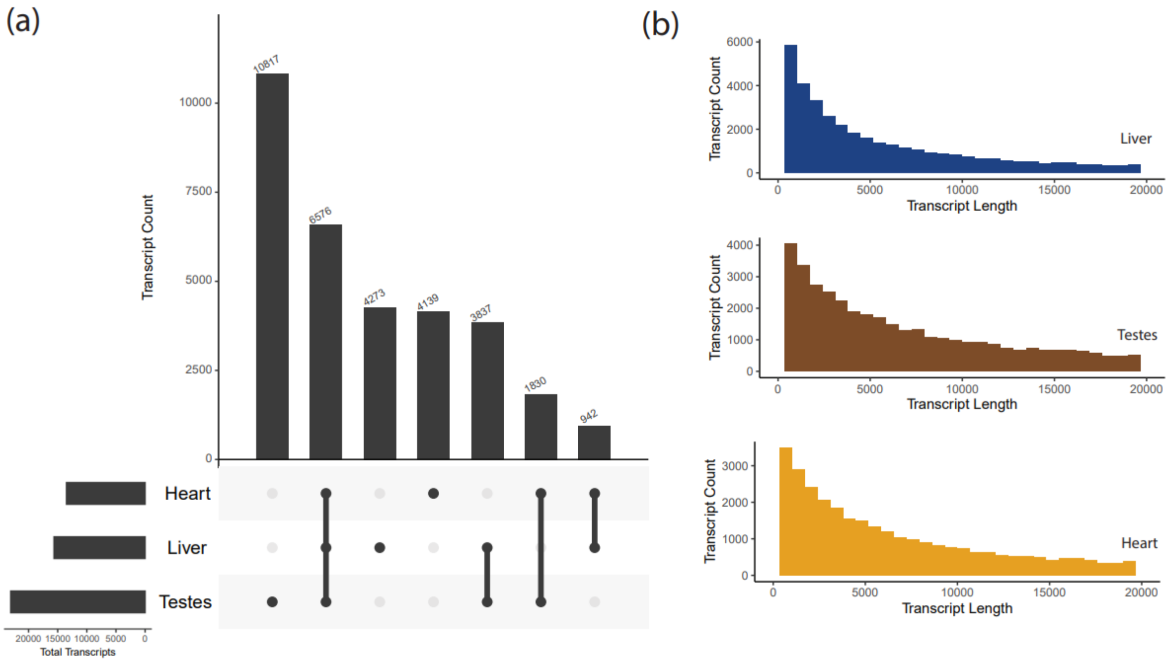


**Figure S11 | *Acridotheres tristis* transcriptome summary information**. Panel **(a)** depicts overlap between tissue-specific transcripts assembled by STRINGTIE v2.2.20, calculated using GFFCOMPARE v0.12.6 and visualised in R using the package UPSETR. Panel **(b)** depicts size classes of transcripts from each tissue.

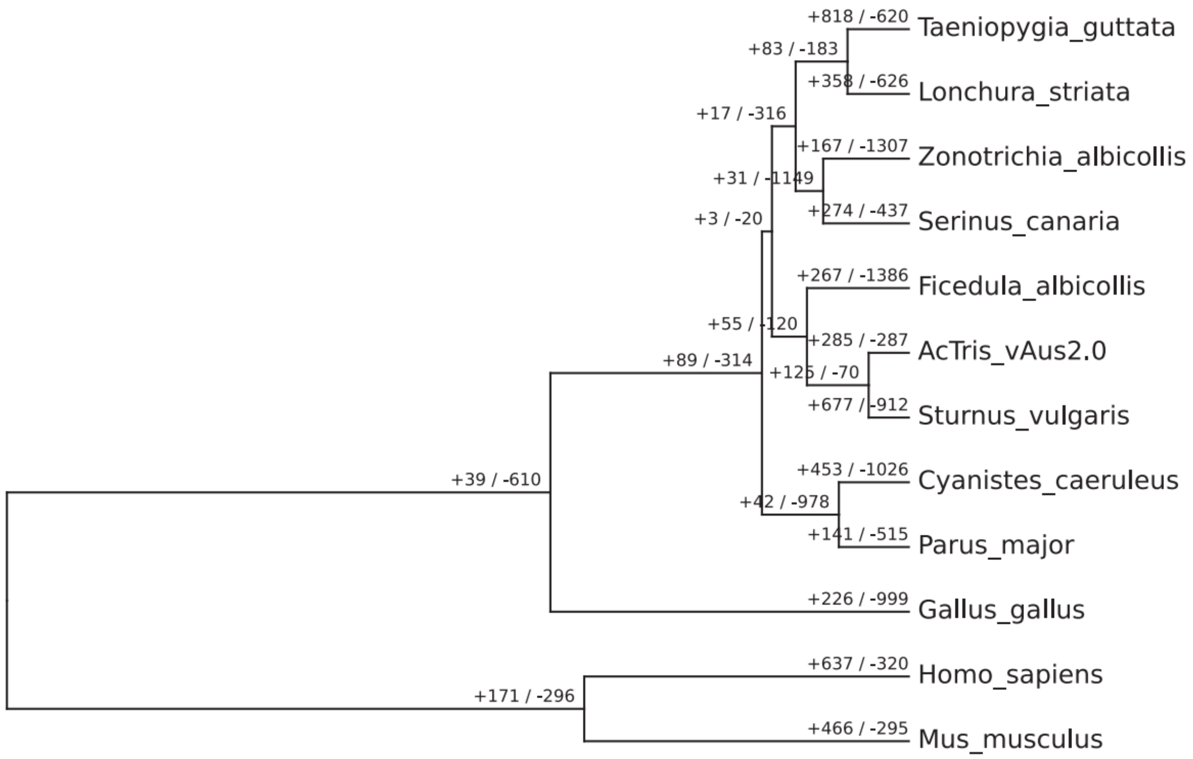


**Figure S12 | All expanded and contracted phylogenetic hierarchical orthogroups** across the 12 species analysed with ORTHOFINDER and CAFE5.

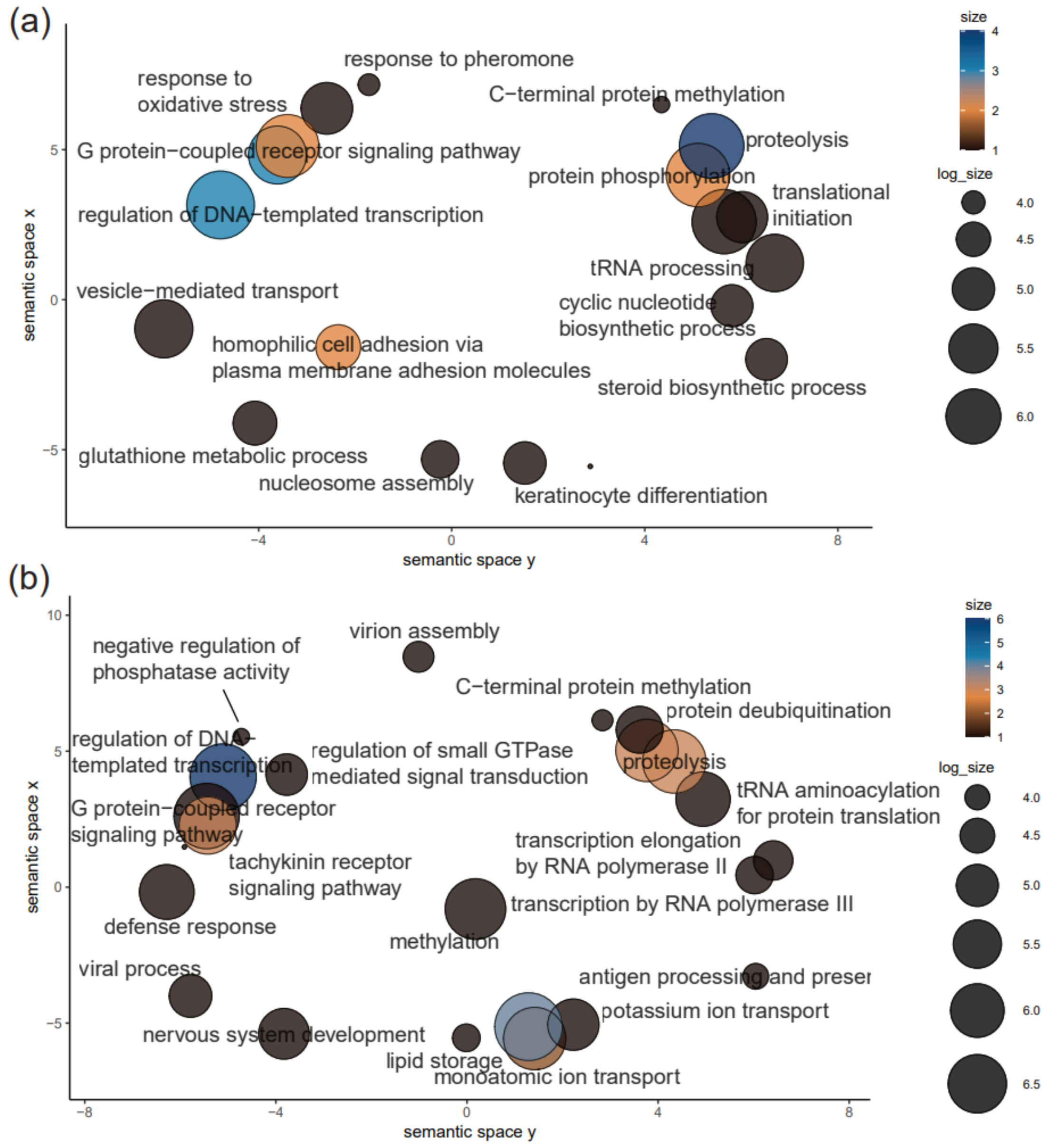


**Figure S13 | REVIGO summary of the biological process captured in gene ontology terms** associated with orthogroups determined to be to be under panel (a) significant expansion and panel (b) significant contractions in *A. tristis.* Colour represents across how many separate significantly expanded or contracted orthogroups the GO term was independently annotated using INTERPROSCAN, while log size represents representing the weighting of that GO term in the dataset once redundant GO terms were collapsed during REVIGO analysis and visualisation. Therefore, both colour and circle size capture the level of representation of the term but do so at different steps of the analysis. Biological processes are written in grey and black for legibility reasons.

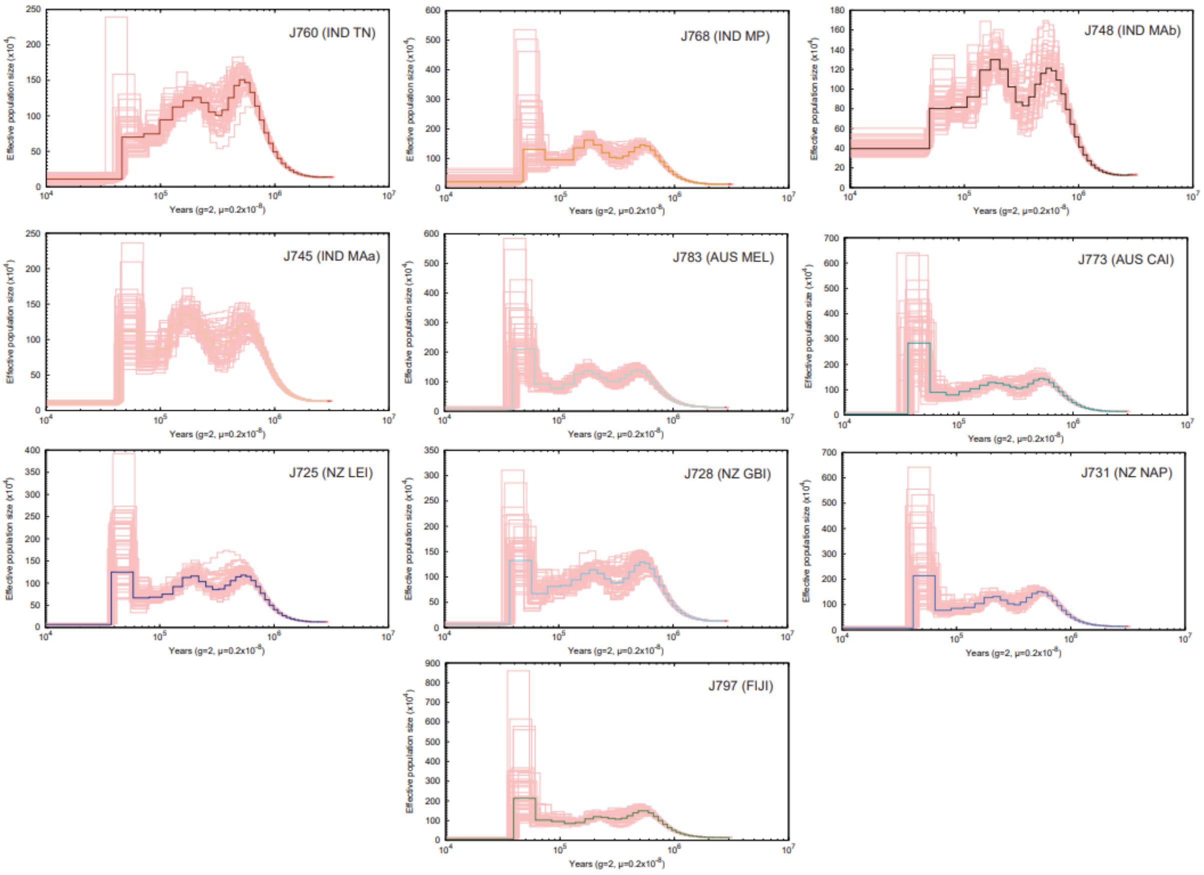


**Figure S14 | PSMC bootstrapping plots** with colour scheme matching Fig. 6 from the main manuscript indicated next to each plot.

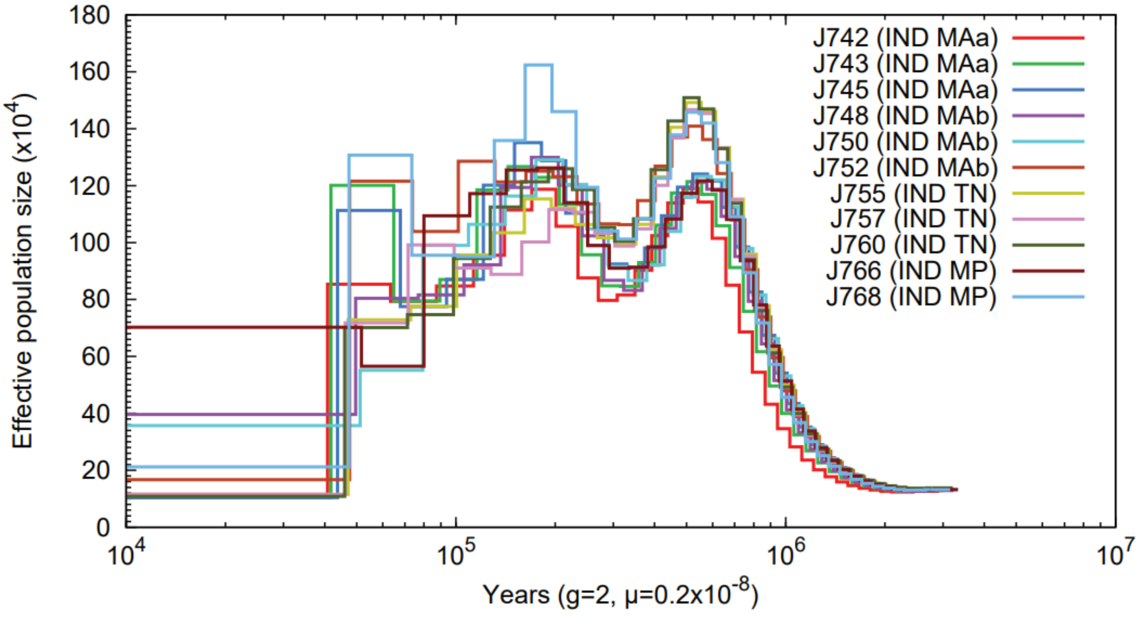


**Figure S15 | PSMC plots** for native range individuals.
